# The Wernicke conundrum revisited: evidence from connectome-based lesion-symptom mapping

**DOI:** 10.1101/2021.10.25.465746

**Authors:** William Matchin, Dirk-Bart den Ouden, Gregory Hickok, Argye E. Hillis, Leonardo Bonilha, Julius Fridriksson

**Author notes:** Corresponding author, 915 Greene St., Discovery 1, Room 202D, Columbia, SC 29208, USA.

## Abstract

Wernicke’s area has been assumed since the 1800s to be the primary region supporting word and sentence comprehension. However, Mesulam et al. (2015; 2019) raised what they termed the ‘Wernicke conundrum’, noting widespread variability in the anatomical definition of this area and presenting data from primary progressive aphasia that challenged this classical assumption. To resolve the conundrum, they posited a ‘double disconnection’ hypothesis: that word and sentence comprehension deficits in stroke-based aphasia result from disconnection of anterior temporal and inferior frontal regions from other parts of the brain due to white matter damage, rather than dysfunction of Wernicke’s area itself. To test this hypothesis, we performed lesion-deficit correlations, including connectome-based lesion-symptom mapping, in four large, partially overlapping groups of English-speaking chronic left hemisphere stroke survivors. After removing variance due to object recognition and associative semantic processing, the same middle and posterior temporal lobe regions were implicated in both word comprehension deficits and complex noncanonical sentence comprehension deficits. Connectome lesion-symptom mapping revealed similar temporal-occipital white matter disconnections for impaired word and noncanonical sentence comprehension, including the temporal pole. We found an additional significant temporal-parietal disconnection for noncanonical sentence comprehension deficits, which may indicate a role for phonological working memory in processing complex syntax, but no significant frontal disconnections. Moreover, damage to these middle-posterior temporal lobe regions was associated with both word and noncanonical sentence comprehension deficits even when accounting for variance due to the strongest anterior temporal and inferior frontal white matter disconnections, respectively. Our results largely agree with the classical notion that Wernicke’s area, defined here as middle superior temporal gyrus and middle-posterior superior temporal sulcus, supports both word and sentence comprehension, suggest a supporting role for temporal pole in both word and sentence comprehension, and speak against the hypothesis that comprehension deficits in Wernicke’s aphasia result from double disconnection.

## 1 Introduction

The classical model of language in the brain posits a primary role for Wernicke’s area, roughly thought to be the posterior superior temporal lobe and inferior parietal lobe (with definitions varying widely)^1–3^, in both word and sentence comprehension^4–7^, defined as a set of processes involving the transformation of an auditory signal onto linguistic representations at the lexical and syntactic levels^8^. The major motivation for the primacy of Wernicke’s area in word and sentence comprehension is the notable comprehension deficits in Wernicke’s aphasia, typically involving posterior temporal-parietal damage^9,10^, which roughly corresponds to the region classically associated with Wernicke’s area^2,3^.

Recently, the classical view has been questioned from the perspective of primary progressive aphasia (PPA). Mesulam et al.^1^ analyzed patterns of cortical thinning in 72 people with PPA on word and sentence comprehension. Their sentence comprehension task minimized lexical access demands by using sentences with highly frequent words (e.g., boy, girl, dog, cat, kiss, chase) and noncanonical structures (object-first sentences such as passives, which do not conform to expected word order), to emphasize grammatical processing. By contrast, their word comprehension task (PPVT) maximized lexical-semantic demands, and the variance due to object recognition and non-verbal semantics was factored out using the Pyramids and Palm Trees test^11^ (PPT) as a covariate. They found no association of posterior temporal-parietal atrophy with word comprehension deficits and minimal association of atrophy in this region with deficits in noncanonical sentence comprehension. By contrast, atrophy of the anterior temporal lobe (ATL), primarily the temporal pole, was strongly associated with word comprehension (but not sentence comprehension) impairment, whereas inferior parietal and frontal degeneration was associated with sentence comprehension (but not word comprehension) impairment. Mesulam et al.^12^ followed up this study by showing that repetition (but not word comprehension) was associated with degeneration of superior posterior temporal and inferior parietal lobes. Overall, they associated the temporal-parietal territory commonly attributed to Wernicke’s area with a phonological working memory function, not critical for basic aspects of linguistic processing.

To explain the discrepancy of their results with the literature on Wernicke’s aphasia, Mesulam et al.^1^ posited a “double disconnection” hypothesis. Given that strokes often impinge on white matter, and that disconnection due to white matter damage can impair language abilities independently of cortical grey matter damage^13,14^, they suggested that lesions to temporal-parietal cortex may impair both word and sentence comprehension, not through damage to the relevant cortical centers themselves but through disconnection^15^. In light of their cortical thinning results in PPA, they proposed that the ATL is the key region underlying word comprehension and that frontal cortex (primarily Broca’s area or the posterior two thirds of the inferior frontal gyrus, IFG) is the key region for sentence/syntactic processing, both of which may be disconnected due to posterior temporal-parietal strokes.

There are two problems with this logic. First, functional neuroimaging studies have shown that the spatial location and extent of activation of syntactic processing and lexical access, the processes leading up to and including word retrieval (but not conceptual retrieval), are remarkably similar (for meta-analyses and reviews, compare Hagoort & Indefrey^16^ for syntactic processing and Lau et al.^17^ for lexical access). This reliably includes the posterior superior temporal sulcus (STS) and posterior middle temporal gyrus (MTG), overlapping with the traditional Wernicke’s territory (although somewhat more anterior than traditionally depicted). Other regions, including prominently the IFG, ATL, and inferior angular gyrus have also been regularly implicated in lexical-semantic processing broadly construed^18,19^. Regardless of the explanation for this overlap of lexical and syntactic processing in the middle-posterior temporal lobe^20,21^, and whether or not the middle-posterior temporal lobe is selectively involved in comprehension-related processes, these activations should be accounted for via some mechanism, and suggest (though do not prove) a functional role in comprehension.

Second, a large body of lesion-symptom mapping (LSM) studies in post-stroke aphasia have shown that damage to Broca’s area and surrounding areas is not reliably implicated in sentence or syntactic comprehension deficits^22–27^ There may be a minor supporting role for frontal cortex in supporting the processing of particularly difficult sentence structures^28^, although the location of the lesion correlates of such deficits is inconsistent^22,29–32^ (for a review, see Matchin & Hickok^21^). If noncanonical sentence comprehension deficits in Wernicke’s aphasia are primarily due to disconnection of Broca’s area/posterior IFG from the rest of the language network, then one would expect frontal lesions to also impair syntactic comprehension. However, this is not a pattern that is reliably seen, in contrast to that associated with posterior temporal-parietal damage, which reliably predicts sentence comprehension impairments^22,23,25,26,30,31,33^.

For these reasons, the dissociation, both in behavior and lesion correlates, between word and sentence comprehension in PPA reported by Mesulam et al.^1^ is surprising and deserves scrutiny. We examined these issues, testing the hypothesis of Mesulam et al. that overlap of word and sentence comprehension deficits in post-stroke aphasia is due to distinct disconnection patterns. We analysed data in a large group of people with post-stroke aphasia using LSM as well as connectome-based lesion-symptom mapping (CLSM), which uses diffusion tensor imaging to ascertain the strength of white matter connections between regions associated with behavioral scores^34–36^. In a previous report^34^, our research group performed LSM and CLSM analyses of word comprehension alone (with PPT as a covariate) in a smaller group of subjects (43), finding that impaired word comprehension was associated with damage to the inferior temporal gyrus and disconnection between posterior middle temporal gyrus and inferior temporal gyrus. A follow-up study^8^ expanded the number of subjects (99) and performed LSM analyses, finding the most robust association of word-level deficits (with PPT as a covariate) with damage to middle-posterior superior temporal gyrus (STG) and STS.

Here we expand the number of subjects further and add several behavioral measures to directly compare our results to Mesulam et al., crucially including a measure of noncanonical sentence comprehension and corresponding CLSM analyses. We predicted that both word and noncanonical sentence comprehension would primarily involve damage to the middle-posterior temporal lobe and similar disconnection patterns throughout the temporal lobe, and that damage to these areas would predict comprehension impairments above and beyond the contribution of the relevant disconnection patterns. We expected a possible implication of parietal damage and/or disconnection in noncanonical sentence comprehension due to phonological working memory demands that are strongly associated with parietal areas^37–40^.

## 2 Materials & Methods

### 2.1 Subjects and measures

We performed behavioral and lesion mapping analyses in four partially overlapping groups of English-speaking subjects (Table 1). We collected anatomical images and created lesion maps for all subjects, but diffusion tensor data for our CLSM analyses was unavailable for several subjects. Group 1 consisted of 218 subjects (199 included in CLSM) who were assessed on the Western Aphasia Battery-Revised^41^, with subsets of this group being assessed on a variety of other measures. Group 2 consisted of a subset of 180 subjects (167 included in CLSM) who were assessed on the Pyramids and Palm Trees test^11^. Group 3 consisted of a subset of 130 subjects (127 included in CLSM) who were assessed on one of two similar tests of sentence comprehension ability involving picture-matching and a variety of semantically reversible canonical and noncanonical sentence structures, the Sentence Comprehension Test subset of the Northwestern Assessment of Verbs and Sentences (NAVS)^42^ (N = 82) or a task which we label here the Icelandic Task (since its translation was first used in a sample of Icelandic stroke survivors with aphasia^31,43^) (N = 48). Finally, Group 4 consisted of a subset of 92 subjects (90 included in CLSM) who were evaluated for expressive agrammatism using consensus perceptual ratings of spontaneous speech samples in one of two studies, Den Ouden et al.^44^ (N = 39) or Matchin et al.^45^ (N = 53).

**Table 1.** Subject demographic data.

|  | **Group 1** | **Group 2** | **Group 3** | **Group 4** |
| --- | --- | --- | --- | --- |
| **Measures** | Western Aphasia Battery-Revised | Pyramids and Palm Trees | Noncanonical Sentence Comprehension | Perceptual Ratings of Expressive Agrammatism |
| **Total number of subjects** | N = 218 (199 included in CLSM) | N = 180 (167 included in CLSM) | Total N = 130 (127 included in CLSM)<br>Icelandic task: N = 48<br>NAVS: N = 82 | Total N = 92 (90 included in CLSM)<br>Den Ouden et al. (2019): N = 39<br>Matchin et al. (2020): N = 53 |
| **Sex** | 133 Male, 85 Female | 112 male, 68 female | 83 male, 47 female | 60 male, 32 female |
| **Mean age at testing (years)** | 60.3 (SD = 11.4) | 60.5 (SD = 11.4) | 60.0 (SD = 10.7) | 58.7 (SD = 11.3) |
| **Mean months post-first stroke at initial testing** | 42.6 (SD = 48.3) | 41.2 (SD = 47.6) | 45.3 (SD = 50.4) | 45.6 (SD = 48.5) |
| **Mean number of** | 1.2 (SD = 0.5) | 1.2 (SD = .5) | 1.2 (SD = 0.5) | 1.1 (SD = 0.4) |
| **strokes** |  |  |  |  |
| **Mean education (years)** | 15.0 (SD = 2.3), *N = 210 | 15.0 (SD = 2.3), *N = 175 | 15.4 (SD = 2.4), *N = 128 | 15.3 (SD = 2.4) |
| **Mean lesion volume (mm<sup>3</sup>)** | 120,855 (SD = 97,488) | 128,446 (SD = 97,036) | 111,267 (SD = 92,645) | 96,175 (SD = 84,514) |
| **Mean WAB-R AQ** | 61.4 (SD = 28.1) | 60.0 (SD = 26.6) | 65.3 (SD = 26.9) | 75.8 (SD = 19.9) |
| **Mean WAB-R Auditory Word Recognition** | 50.0 (SD = 13.2) | 49.8 (SD = 12.9) | 51.0 (SD = 12.7) | 54.7 (SD = 9.0) |
| **Mean WAB-R Sequential Commands** | 51.4 (SD = 23.5) | 50.4 (SD = 22.5) | 54.5 (SD = 22.7) | 61.0 (SD = 19.5) |
| **Mean WAB-R Repetition** | 5.5 (SD = 3.4) | 5.3 (SD = 3.3) | 5.9 (SD = 3.3) | 7.0 (SD = 2.7) |

SD = standard deviation. AQ = aphasia quotient of the WAB-R, a summary measure of overall language ability; WAB-R = Western Aphasia Battery-Revised; SD = standard deviation. \*education information was not available for all subjects, the number for which education information indicated here.

All subjects provided informed consent to participate in this study, which was approved by the Institutional Review Boards at the University of South Carolina and the Medical University of South Carolina. All subjects were native speakers of American English and had at least one ischemic stroke to the left hemisphere at least six months prior to study inclusion.

In order to compare our study effectively with the studies of Mesulam et al. in PPA^1,12^, we included a similar set of six behavioral measures: the WAB-R Auditory Word Comprehension subtest, the Pyramids and Palm Trees Test (PPT), the Philadelphia Naming Test, Noncanonical Sentence Comprehension, the WAB-R Repetition subtest, and our perceptual ratings of Expressive Agrammatism. Each of these measures is described below. Out of these six measures, we ultimately performed four lesion mapping analyses, described in section 2.3.

The Auditory Word Recognition subtest of the WAB-R^41^ involves asking the subject to point to real-world objects or printed images (n = 60). Importantly, the test does not require syntactic parsing in order to perform correctly, as the subject only needs to identify the lexical item presented in each item. Our *Word Comprehension* measure used for lesion analyses consisted of this measure incorporating the Pyramids and Palm Trees (PPT) measure described below as a covariate.

The WAB-R Repetition subtest^41^ (referred to subsequently as *Repetition*) involves presenting a series of increasingly complex utterances and requiring subjects to repeat them verbatim. Scoring is based on the number of words correctly recalled in the correct order.

The Pyramids and Palm Trees test (PPT)^11^ involves presenting a target picture with two candidate pictures below that are possible associates of the target picture (e.g., the target picture could be a pyramid, with candidate pictures of palm trees and pine trees). The subject is required to point to the candidate picture that is more related to the target picture (e.g. the palm trees). There are a total of 52 trials. This task was included in order to provide a control for object recognition and non-verbal semantic processing in the WAB-R word comprehension test (as in Mesulam et al.^1^ and Bonilha et al.^34^).

The Philadelphia Naming Test (PNT)^46^ is a 175-item assessment of picture naming, involving the presentation of a number of simple line drawings. Here we used the total number of items correctly named on the task (rather than phonological or semantic errors).

Our *Noncanonical Sentence Comprehension* measure was derived from either the NAVS (N = 82) or the Icelandic task (N = 48). The NAVS Sentence Comprehension Test^42^ involves testing the comprehension of a variety of canonical and noncanonical sentence types, each with five total trials, assessed via pointing to the correct picture. Mesulam et al. (2015) assessed the comprehension performance on the three noncanonical sentence types: passives (with a by-phrase) e.g. *the dog is chased by the cat,* object-extracted WH-questions, e.g. *who is the cat chasing?,* and object-relatives, e.g. *Pete saw the boy who the girl is pulling,* for a maximum score of 15. The Icelandic task includes a similar set of canonical and noncanonical sentence types, each with five total trials: passives (with a by-phrase), e.g. *the boy is painted by the girl,* object-extracted WH-questions, e.g. *which boy is the girl painting?*, and object clefts, e.g. *it is the girl that the boy paints.* The sentence types across the two tasks are not strictly identical, but involve essentially the same structures with the same degree of complexity, including the key factor of noncanonical object-first word order. Therefore, for subjects who were not assessed with the NAVS, we calculated the equivalent scores on the Icelandic task (correct noncanonical trials, out of 15 points).

The *Expressive Agrammatism* measure was a perceptual measure of grammatical deficits in speech production. We derived this measure from samples of connected speech production elicited either by describing the Cookie Theft picture^47^ (N=39) as reported in Den Ouden et al.^44^ or retelling the story of Cinderella in their own words^48^ (N=53), as reported in Matchin et al.^45^. Production samples were rated independently by speech and language experts for the systematic simplification of sentence structure and omission of function words and morphemes. This resulted in a categorical assessment for each subject, either agrammatic or not.

### 2.2 Brain imaging and lesion mapping

We acquired anatomical MRI and DTI data using the same parameters and procedures as described in previous studies^30,34,44^. High-resolution neuroimaging data (T1- and T2-weighted images) were collected at the University of South Carolina and the Medical University of South Carolina on a 3T Siemens Trio scanner with a 12-element head coil. Lesions were demarcated onto each subject’s T2 image by an expert neurologist (Dr. Leonardo Bonilha) or an expert cognitive neuroscientist (Dr. Roger Newman-Norlund) extensively trained by Dr. Bonilha (with consultation as needed with an expert on lesion mapping, Dr. Chris Rorden), both blind to the behavioral data. Lesion maps were then aligned to the high resolution T1 image. Lesions were replaced with the corresponding brain structure from the intact hemisphere, and this image as well as the lesion map in subject space were subsequently warped to MNI space using SPM12^49^. The warped lesion map was then binarized with a 50% probability threshold, which was used to perform voxel-wise and region of interest (ROI) analyses.

Lesion overlap maps for both groups are shown in Figure 1. Overall, there was good coverage in perisylvian cortex, covering all relevant language-related regions (see lesion overlap maps in Figure 1). We used the JHU atlas^50^ (depicted in Supplementary Figure 2, full list of relevant region abbreviation definitions in Supplementary Table 1) for lesion-symptom mapping (LSM) and connectome-based lesion-symptom mapping (CLSM) analyses. The parcellations of the JHU atlas provide good alignment with language-relevant brain regions and have roughly equivalent parcel sizes relative to parcellations included in other atlases, such as the AICHA atlas^51^. The JHU atlas contains two ROIs containing the polar and lateral anterior components of the superior temporal gyrus (STG) and middle temporal gyrus (MTG), the STG pole and MTG pole, which correspond to key components of the ATL. We note that Mesulam et al.^1,12^ define the ATL as including regions more inferior and posterior to these two ROIs. However, we also note that Mesulam et al.^1^ attributed a dominant role to the polar part of the ATL, p. 2431: “...the ATL peak atrophy sites associated with severe word comprehension impairment consistently included the temporal pole...ATL neuronal loss causes major word comprehension impairments only if it extends anteriorly all the way into the polar region”, thus we consider these two ROIs as well-defined to test the disconnection claim. However, as described below in section 2.3.4, we accommodated the possibility that more posterior portions of the ATL might be relevant to the disconnection claims in our combined LSM and CLSM analyses.

**Figure 1.**
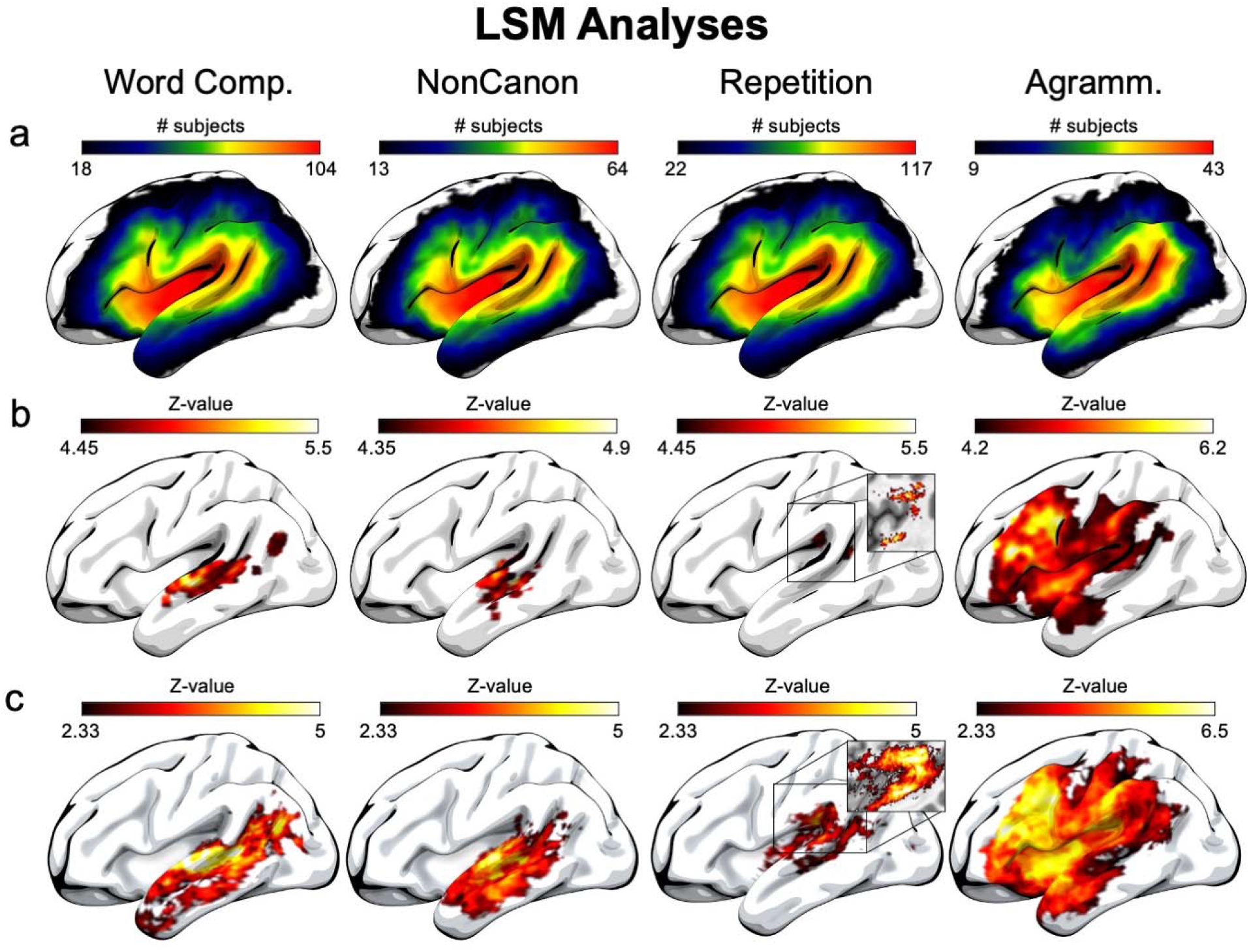
Lesion-symptom mapping (LSM) analyses. a) Lesion overlap maps for each group associated with the behavioral measures for which we analyzed lesion-deficit correlations. Note that the lower value in each color spectrum indicates the minimum number of subjects with damage to each region that was required for our 10% lesion load threshold. b) Voxel-wise univariate LSM analyses for each behavioral measure (incorporating lesion volume as a covariate), with a permutation correction for multiple comparisons (10,000 permutations, corrected p < 0.05), with results spatially smoothed for improved visibility. c) Uncorrected (voxel-wise p < 0.01) and unsmoothed voxel-wise univariate LSM analyses for each behavioral measure (incorporating lesion volume as a covariate). Insets show white matter damage in a medial slice of the volumetric data underlying the cortical surface not adequately depicted in the surface rendering. Comp = Comprehension, NonCanon = Noncanonical Sentence Comprehension, Agramm. = Expressive Agrammatism.

### 2.3 Analyses

#### 2.3.1 Behavioral

To examine the relationship among the six total behavioral variables, we performed nonparametric Spearman correlations using JASP^52^. Missing data were accounted for with pairwise deletion. Non-parametric Spearman correlations were used given that the behavioral variables did not have pairwise normal distributions. Lesion volume is strongly correlated with aphasia severity in chronic stroke^53–55^, including in our data (one-tailed Spearman’s rho correlating WAB-R AQ and lesion volume = −0.694, p = 6.254 x 10^-33^), and therefore a potential confound^56,57^, which we controlled by using lesion volume as a covariate in our LSM analyses. Similarly, we report behavioral correlations both with and without lesion volume as a covariate to evaluate overall severity effects. Mesulam et al.^1^ did not correct their behavioral correlation analyses for multiple comparisons, but given the large number of behavioral correlations we performed (15 pairwise correlations without lesion volume as a covariate, 15 pairwise correlations with lesion volume as a covariate), we report whether or not each correlation survived a Bonferroni correction for multiple comparisons, treating each set of 15 correlations (with and without the lesion volume covariate) as a separate family of tests, using an adjusted alpha level of p < 0.003.

#### 2.3.2 LSM

With respect to LSM, using NiiStat (https://www.nitrc.org/projects/niistat/) we performed both voxel-based and ROI-based analyses relating each of the four selected behavioral variables to lesion location. ROI analyses were included in order to maximize statistical power and afford an opportunity to combine the LSM and CLSM data to test the hypothesis of Mesulam et al. that disconnection accounts for the substantial association of damage to middle-posterior temporal lobe regions and comprehension impairments. For voxel-based analyses, we performed one-tailed t-tests within each voxel comparing the magnitude of the behavioral measure for subjects with and without damage to that voxel (results were converted to Z-values for ease of interpretation). For ROI analyses, we performed univariate linear regression analyses relating proportion damage within the grey matter of each parcellated region (the number of voxels that were damaged divided by the total number of voxels in each region) contained within the JHU atlas (depicted in Supplementary Figure 2), except for Expressive Agrammatism, which was logistic regression^50^. Both voxel-based and ROI-based analyses were only performed within voxels/regions that had at least 10% of subjects with damage located there^57,58^ and were corrected for multiple comparisons using permutation tests (10,000 permutations) with a corrected alpha threshold of p < 0.05. We performed analyses for four behavioral measures that provided maximum comparison to the analyses of Mesulam et al.^1,12^: WAB-R Auditory Word Recognition with PPT as a covariate *(Word Comprehension), Noncanonical Sentence Comprehension,* WAB-R Repetition *(Repetition),* and *Expressive Agrammatism.* Lesion volume was included as a covariate in all LSM analyses to ensure accurate localization^56,57^ (we also report correlations between each major behavioral measure and overall lesion volume in order to assess the likelihood of Type 2 errors of including lesion volume as a covariate, that is, of overly conversative LSM analyses resulting from the lesion volume covariate). Univariate analyses were performed as opposed to multivariate approaches such as Support Vector Regression (SVR) in order to match the univariate analyses of Mesulam et al.^1^, for straightforward interpretation of statistical results, and because univariate approaches have been shown to outperform multivariate approaches with large sample sizes and adequate lesion load restrictions^57^. To supplement our voxel-based analyses, we also report reduced threshold (voxel-wise p < 0.01) lesion maps for all behavioral variables in order to provide the fullest picture of the lesion data without obscuring near-threshold results.

To address the question of regions that cause both word and sentence comprehension impairment, we calculated the overlap of the corrected voxel-based lesion maps for the Word Comprehension and Noncanonical Sentence Comprehension measures. Regions with significant overlap could be considered candidates for an updated anatomical definition of “Wernicke’s Area” according to its functional definition as causing both of these impairments.

Finally, an additional issue concerns the fact that our Word Comprehension and Noncanonical Sentence Comprehension measures do not control for prelexical speech perception impairments. We performed analyses of two additional measures including the Repetition score as an additional covariate to attempt to control for these impairments. We first combined the behavioral scores with these covariates in a linear regression model (WAB-R Auditory Word Recognition with both PPT and Repetition, and Noncanonical Sentence Comprehension with Repetition), saved the residual values, and then performed LSM analyses of the two resulting measures in the same manner as described above.

#### 2.3.3 CLSM

For CLSM, we analyzed the diffusion-weighted images that were acquired for each subject and estimated the pairwise connection strength between all regions within the JHU atlas, including both the left and right hemispheres^50^ (see Supplementary Figure 2 for a depiction of a subset of the regions contained within the JHU atlas). First, each subject’s T1 image was used to register white matter and region parcellation to the diffusion-weighted images. Next, we estimated connection strength between regions using fiber count (corrected for distance and region volume) for each of the pairwise connections by using probabilistic tractography as implemented in FSL’s FDT method^59^, using an approach that minimizes the potential distorting effects of brain damage on fiber tracking^13^. Pairwise connectivity was calculated as the fiber count arriving in one region when another region was seeded and averaged with the fiber count calculated in the reverse direction. The lesioned tissue was removed from all tractography tracings in order to maximize accuracy, which also minimizes the effect of lesion volume on the final analyses. The estimated number of streamlines from a completely destroyed region was set to zero. Overall, the number of streamlines that can be estimated between a pair of regions is constrained but not determined by the amount of damage to each region, as the damage to intervening white matter pathways is crucial to estimating the intact connections between these regions. The number of estimated tracts connecting pairs of regions was divided by the total volume of both regions, controlling for unequal region sizes. For full details of the methodological approach, see other publications from our research group using the same method^34,36^.

We then performed linear regression analyses relating scores on each behavioral measure with the estimated connection strength for each connection using NiiStat (https://www.nitrc.org/projects/niistat/), correcting for multiple comparisons using permutation tests (10,000 permutations) and a corrected alpha threshold of p < 0.05 for each of the four behavioral measures (Word Comprehension, Noncanonical Sentence Comprehension, Repetition, and Expressive Agrammatism). When including lesion volume as a covariate, we identified very few significant disconnections associated with noncanonical sentence comprehension deficits and no significant disconnections associated with any of the other three behavioral measures we assessed (Word Comprehension, Repetition, and Expressive Agrammatism). It is as yet unclear whether lesion volume is critical for accurate localization as in LSM^56,57^, although as noted above, lesion volume was already factored into our analyses by removing damaged tissue from estimated tractographies. Therefore, we focus our results on analyses that did not include lesion volume as a covariate in our CLSM analyses, in line with previous reports from our research group^34,44^.

#### 2.3.4. LSM and CLSM combined

The ‘double disconnection’ hypothesis of Mesulam et al. entails that the association between damage to temporal-parietal regions (roughly Wernicke’s area) with word and sentence comprehension deficits in Wernicke’s aphasia can be explained via disrupted connections due to damage to underlying white matter between the ATL and frontal cortex to potentially various other regions of the brain (Mesulam, personal communication). If this were true, then there should be no robust independent contribution of damage to these regions above and beyond the extent of disruption to these connections. To test this, we combined the LSM and CLSM data by assessing whether lesion-deficit correlations for these behavioral measures would still be statistically robust when incorporating connection strength as a covariate.

First, we identified the relevant regions associated with Word Comprehension and Noncanonical Sentence Comprehension using the ROI-based LSM analyses described in section 2.3.2. Both measures identified the same set of regions: superior temporal gyrus, middle portion (STG), posterior superior temporal gyrus (pSTG), middle temporal gyrus, middle portion (MTG), and posterior middle temporal gyrus (pMTG). For Word Comprehension, for the more posterior of these regions (pSTG and pMTG) we identified the single strongest disconnection involving a more anterior temporal lobe region associated with behavioral deficits, MTG ↔□ superior occipital gyrus (SOG) (Z = 4.04). We then did the same for the middle temporal regions, STG and MTG, identifying the MTG pole SOG (Z = 3.98). For Noncanonical Sentence Comprehension, we identified the single strongest disconnections involving all four of these regions identified in the ROI-based LSM analyses and a frontal lobe region. Given that no frontal disconnections were identified that survived the multiple comparisons correction in the CLSM analyses, we selected the single strongest sub-threshold frontal disconnection for these regions out of the set of IFG pars opercularis (IFG opercularis), IFG pars triangularis (IFG triangularis), IFG pars orbitalis (IFG orbitalis), and posterior middle frontal gyrus (pMFG), given that the IFG regions are classically associated with grammatical processing and the pMFG was identified in our ROI-based LSM analysis of Expressive Agrammatism. This identified the connection between IFG orbitalis and the Cuneus (Z = 3.66) as the most strongly disconnected. The selected regions and connections for these analyses are depicted in Figure 4.

We then performed linear regression analyses to predict behavioral scores using proportion damage to each ROI, incorporating both lesion volume and the relevant connection for that region as covariates. In this way, we tested whether the strongest anterior temporal disconnection could account for the lesion-deficit correlations in the posterior temporal lobe for Word Comprehension, and whether the strongest frontal disconnection could account for the lesion-deficit correlations in the temporal lobe for Noncanonical Sentence Comprehension. Under the Mesulam et al. hypothesis, we would expect that there would no longer be a significant association between the behavioral scores and damage to middle-posterior temporal lobe regions when these disconnections are considered.

### 2.4 Data Availability

All of the lesion maps and behavioral data for subjects enrolled in this study are available for download at https://www.dropbox.com/sh/3w4aeizgypfs7sd/AAB-W8Yn5qDUFeBj90WKsBqAa?dl=0 (use “wernickeConundrumRevisited_CLSM.xlsx” file).

## 3 Results

### 3.1 Behavioral

Non-parametric Spearman correlations found significant correlations among most of our behavioral variables when lesion volume was not included as a covariate (Table 2, top). In fact, the only correlations that were not significant were between Noncanonical Sentence Comprehension and PPT (although this nearly survived the Bonferroni correction for multiple comparisons), and between Expressive Agrammatism and three measures: Noncanonical Sentence Comprehension, Repetition, and PPT. When lesion volume was included as a covariate, we found overall weaker correlations among the behavioral measures, and Expressive Agrammatism was no longer significantly correlated with any of the other measures (Table 2, bottom).

**Table 2.** Behavioral analyses.

| **Analyses with no covariates** |  |  |  |  |  |
| --- | --- | --- | --- | --- | --- |
|  | PNT | NonCanon | Repetition | PPT | Agramm. |
| Word Comp. | N = 162 | N = 130 | N = 218 | N = 181 | N = 92 |
| | $\rho = 0.565$ | $\rho = 0.415$ | $\rho = 0.738$ | $\rho = 0.622$ | $\rho = 0.377$ |
|  | **$p < 0.001$** | **$p < 0.001$** | **$p < 0.001$** | **$p < 0.001$** | **$p < 0.001$** |
| PNT |  | N = 102 | N = 162 | N = 155 | N = 73 |
| | | $\rho = 0.451$ | $\rho = 0.802$ | $\rho = 0.408$ | $\rho = 0.350$ |
|  |  | **$p < 0.001$** | **$p < 0.001$** | **$p < 0.001$** | **$p < 0.001$** |
| NonCanon |  |  | N = 130 | N = 108 | N = 79 |
| | | | $\rho = 0.441$ | $\rho = 0.236$ | $\rho = 0.101$ |
| | | | **$p < 0.001$** | $p = 0.007$ | $p = 0.197$ |
| Repetition |  |  |  | N = 181 | N = 92 |
| | | | | $\rho = 0.502$ | $\rho = 0.205$ |
| | | | | **$p < 0.001$** | $p = 0.039$ |
| PPT |  |  |  |  | N = 75 |
| | | | | | $\rho = 0.112$ |
| | | | | | $p = 0.164$ |
| **Analyses with lesion volume covariate** |  |  |  |  |  |
|  | PNT | NonCanon | Repetition | PPT | Agramm. |
| Word Comp. | N = 162 | N = 130 | N = 218 | N = 181 | N = 92 |
| | **$\rho = 0.433$** | **$\rho = 0.281$** | **$\rho = 0.549$** | **$\rho = 0.523$** | $\rho = 0.038$ |
| | **$p < 0.001$** | **$p < 0.001$** | **$p < 0.001$** | **$p < 0.001$** | $p = 0.359$ |
| PNT |  | N = 102 | N = 162 | N = 155 | N = 73 |
| | | **$\rho = 0.364$** | **$\rho = 0.756$** | **$\rho = 0.306$** | $\rho = -0.051$ |
| | | **$p < 0.001$** | **$p < 0.001$** | **$p < 0.001$** | $p = 0.683$ |
| NonCanon |  |  | N = 130 | N = 108 | N = 79 |
| | | | **$\rho = 0.317$** | $\rho = 0.146$ | $\rho = -0.137$ |
| | | | **$p < 0.001$** | $p = 0.067$ | $p = 0.875$ |
| Repetition |  |  |  | N = 181 | N = 92 |
| | | | | **$\rho = 0.359$** | $\rho = 0.121$ |
| | | | | **$p < 0.001$** | $p = 0.152$ |
| PPT |  |  |  |  | N = 75 |
| | | | | | $\rho = -0.076$ |
| | | | | | $p = 0.747$ |

Correlation analyses that survived the Bonferroni correction for multiple comparisons within each family of tests (with/without lesion volume covariate) (adjusted alpha of $p < 0.003$ ) indicated in bold. $\rho$ = Spearman's rho, Comp. = Comprehension, PNT = Philadelphia Naming Task, NonCanon = Noncanonical Sentence Comprehension, PPT = Pyramids and Palm Trees, Agramm. = Expressive Agrammatism.

Similar to Mesulam et al.^1^, we found that WAB-R auditory word recognition and PPT scores were strongly correlated, justifying our use of PPT scores as a covariate with WAB-R auditory word recognition to create the Word Comprehension measure, removing variance due to object recognition and non-verbal semantic processing in our lesion-symptom mapping analyses. We also found that Noncanonical Sentence Comprehension and Repetition scores were correlated, suggesting that at least some of the variance in sentence comprehension scores could be due to phonological working memory abilities.

There were notable differences from the behavioral correlations reported by Mesulam et al.^1^. Importantly, we found robust correlations between Noncanonical Sentence Comprehension and WAB-R auditory word recognition, whether or not lesion volume was included as a covariate, unlike Mesulam et al., who found that word and sentence comprehension were not correlated. This is consistent with the classical findings in stroke-based aphasia that word and sentence comprehension deficits coincide. These results underscore the ‘Wernicke Conundrum’: apparent differences in the patterns of word and sentence comprehension deficits between PPA and stroke-based aphasia. We also found that WAB-R Repetition was very robustly correlated with WAB-R auditory word recognition, whereas Mesulam et al. found that word comprehension and repetition were not correlated.

Finally, we also found that Noncanonical Sentence Comprehension was not correlated with Expressive Agrammatism, whether or not lesion volume was included as a covariate. Although correlations between Expressive Agrammatism and other behavioral variables were not assessed in Mesulam et al.^1,12^, this lack of a relationship in our data speaks against one of the major ideas promoted in this work. Namely, they suggested that sentence comprehension deficits in both PPA and stroke-based aphasia are due to degeneration and/or disconnection of a frontal-based grammatical processing center that affects both production and comprehension. Under this perspective, one would expect that both Expressive Agrammatism and Noncanonical Sentence Comprehension would be correlated, but they were not. Note that we did find that Expressive Agrammatism correlated with WAB-R auditory word recognition and picture naming when lesion volume was not included as a covariate; thus the failure to identify a correlation between Expressive Agrammatism and Noncanonical Sentence Comprehension cannot be merely due to a lack of statistical power.

### 3.2 LSM

Voxel-based LSM results are shown in Figure 1 and Supplementary Figure 1. ROI-based LSM results are listed in Table 3. Spearman Correlations between each behavioral measure and overall lesion volume are as follows: Word Comprehension (WAB-R Auditory Word Recognition deficits, with PPT scores as a covariate) and lesion volume, r = −0.354, p = 2.851e-6; Noncanonical Sentence Comprehension and lesion volume, r = −0.314, p = 2.734e-4; Repetition and lesion volume, r = −0.577, p = 9.824e-21; Expressive Agrammatism, r = 0.639, p = 6.965e-12.

**Table 3.**
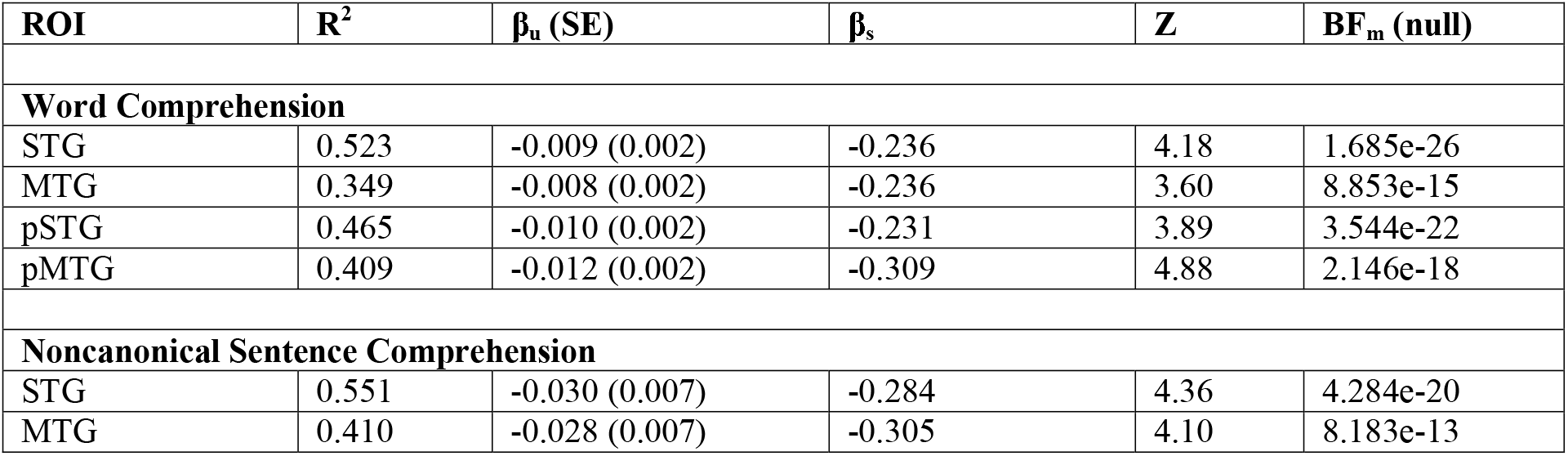
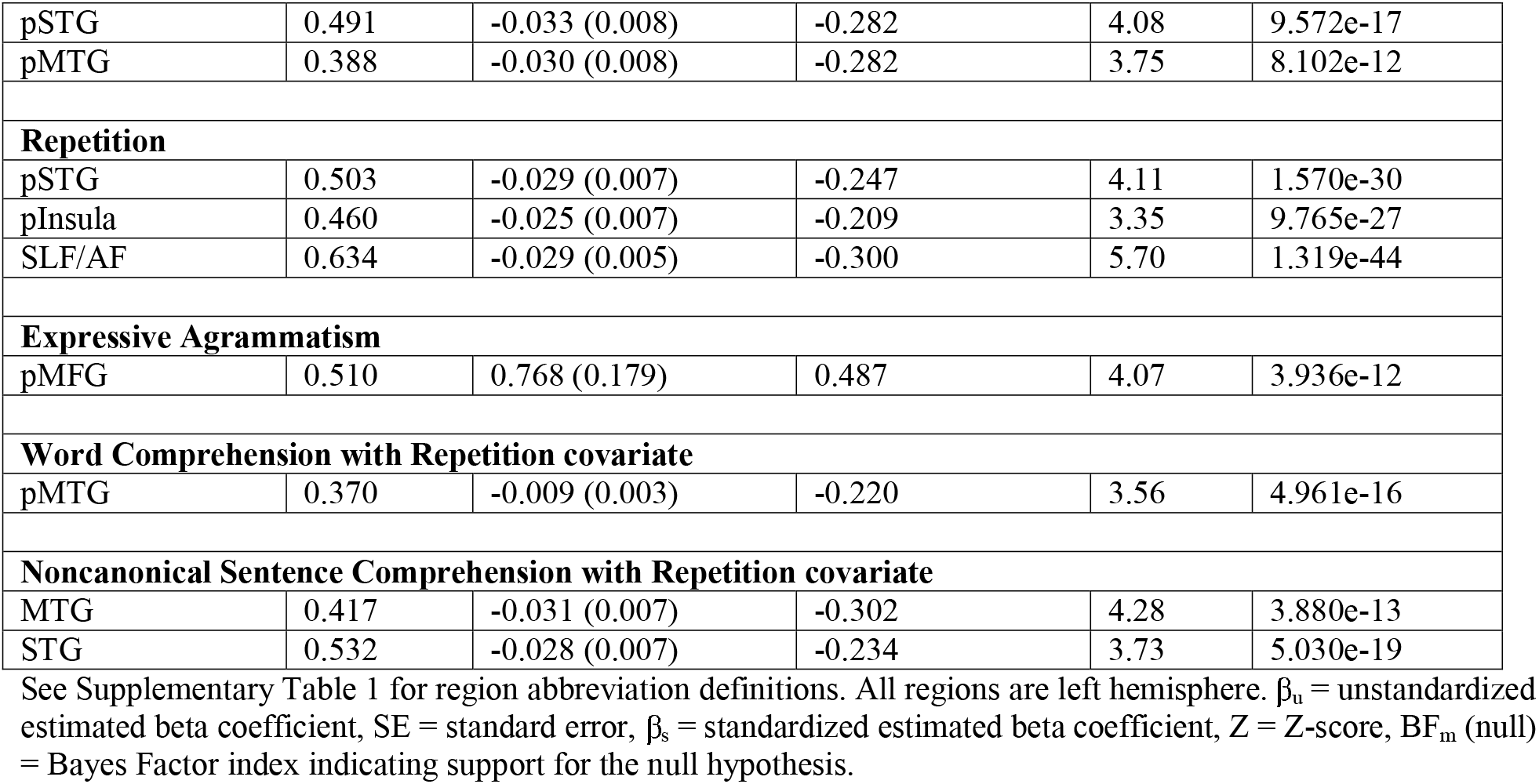
Significant regions from ROI-based LSM analyses.

Focusing on the corrected voxel-based analyses (Figure 1b), Word Comprehension deficits were associated with damage to middle and posterior STG (pSTG) and posterior MTG (pMTG) and to a lesser extent inferior angular gyrus. Noncanonical Sentence Comprehension deficits were associated with a similar lesion distribution, although without including damage to the inferior angular gyrus. The corrected ROI analyses revealed that damage to the same set of four regions was associated with both Word Comprehension and Noncanonical Sentence Comprehension deficits: STG (middle), MTG (middle), pSTG, and pMTG. The supplementary analyses that included Repetition as a covariate for both of these measures revealed largely similar results with substantially reduced statistical strength (Supplementary Figure 1), with a subset of the same regions significantly identified in the ROI analyses (Table 3).

By contrast, the corrected voxel-based LSM analyses revealed that Repetition deficits were primarily associated with damage to posterior STG and superior longitudinal/arcuate fasciculus. The corrected ROI-based analyses revealed similar results, including damage to those regions as well as the posterior insula. Expressive Agrammatism was associated with a widespread pattern of damage in the corrected voxel-based analyses, including most prominently the posterior middle frontal gyrus (pMFG) and superior IFG, although this lesion map extended contiguously into the anterior insula, supramarginal gyrus, inferior angular gyrus, and ATL. The corrected ROI-based analyses of Expressive Agrammatism only revealed significant damage to the pMFG.

The voxel-based overlap analyses of Word Comprehension and Noncanonical Sentence Comprehension (Figure 2) showed substantial overlap of the lesion correlates of these two measures in the middle STG and middle-posterior STS. The centers of mass for the primary clusters of overlapping voxels were located at the following sets of MNI coordinates:

- −48, −34, −1
- −59, −16, 2
- −53, −20, −3.

**Figure 2.**
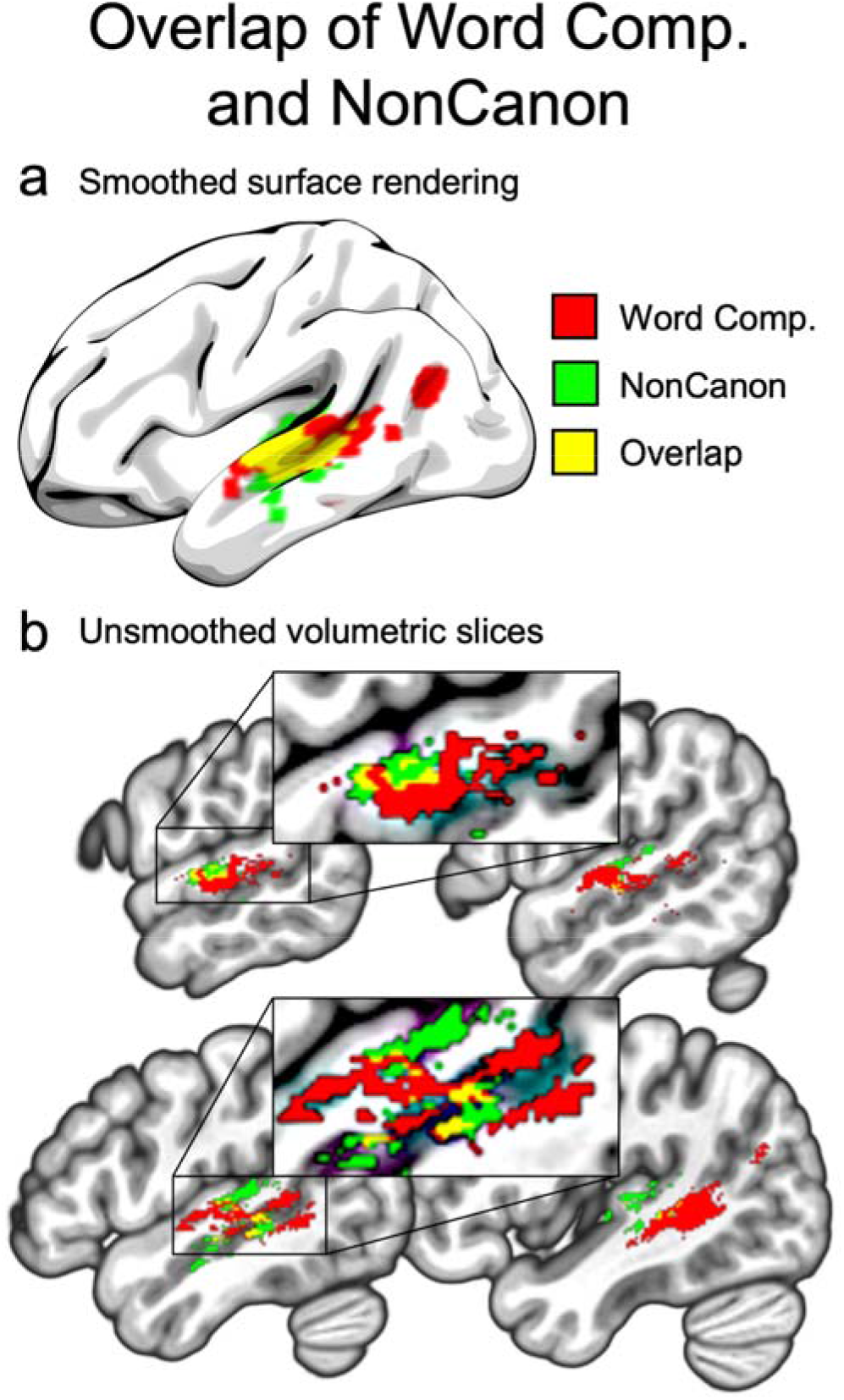
Wernicke’s area localized through the convergence of the lesion maps for word and sentence comprehension deficits. a) Smoothed surface rendering of the overlap (yellow) of the corrected lesion maps for Word Comprehension (red) and Noncanonical Sentence Comprehension (green). b) Unsmoothed slices of the volumetric (non-surface-rendered) overlap. Insets show zoomed-in views of the portions of the slices surrounded by rectangles. Comp = Comprehension, NonCanon = Noncanonical Sentence Comprehension.

This overlap is consistent with the classic picture from stroke-based aphasia in which word and sentence comprehension coincide in Wernicke’s aphasia following damage to posterior temporal lobe, although we note that the strongest overlap occurred in middle STG and posterior-middle STS for both measures. Combined with the robust ROI results in both posterior and middle temporal regions, these results suggest that the middle temporal lobe is crucial to an anatomical definition of Wernicke’s area. The overlap of the supplemental analyses including Repetition as a covariate showed very similar regions but with no strict overlap of Word Comprehension and Noncanonical Sentence Comprehension at the voxel level (Supplementary Figure 1), likely due to the greatly reduced statistical power of these analyses.

### 3.3 CLSM

CLSM results are shown in Figure 3, and the top ten significant disconnections for our primary analyses are listed in Table 4 (the full set of significant disconnections for the CLSM analysis of Repetition are listed in Supplementary Table 3, as are the significant disconnections for our supplementary analyses of Word and Noncanonical Sentence Comprehension including Repetition as a covariate). Both impaired Word Comprehension (WAB-R auditory word recognition with PPT as a covariate) and Noncanonical Sentence Comprehension were associated with several disconnections within the temporal lobe and between temporal and occipital areas, including temporal pole regions. Impaired Noncanonical Sentence Comprehension was associated with one significant disconnection between the superior parietal lobe and the temporal pole, as well as two right hemisphere angular gyrus disconnections involving the left visual cortex. There were no significant frontal disconnections associated with either Word or Noncanonical Sentence Comprehension deficits, and there were no right hemisphere disconnections associated with Word Comprehension deficits. This speaks against the hypothesis of Mesulam et al.^1^ that impaired noncanonical sentence comprehension in stroke-based aphasia results from impaired disconnection of a frontal-based grammatical processing system and that impaired word comprehension might result from disconnection of the left ATL from right hemisphere regions.

**Figure 3.**
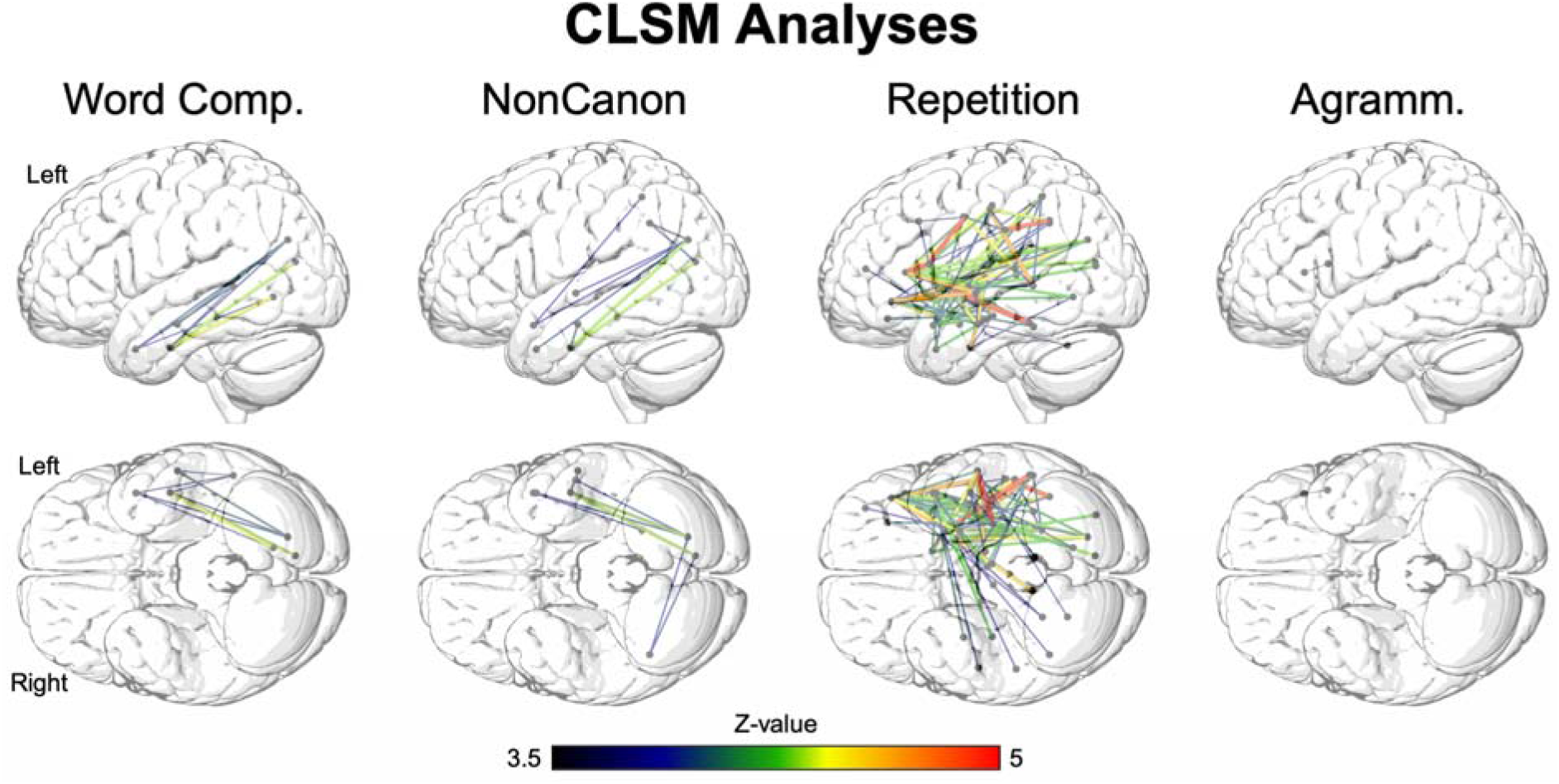
Connectome-based lesion-symptom mapping (CLSM) analyses. Significant connections between ROIs based on the JHU atlas ascertained by diffusion tensor imaging, with a permutation correction for multiple comparisons (10,000 permutations, corrected p < 0.05). Dots indicate regions at the endpoints of significant disconnections. Comp. = Comprehension, NonCanon = Noncanonical Sentence Comprehension, Agramm. = Expressive Agrammatism.

**Table 4.**
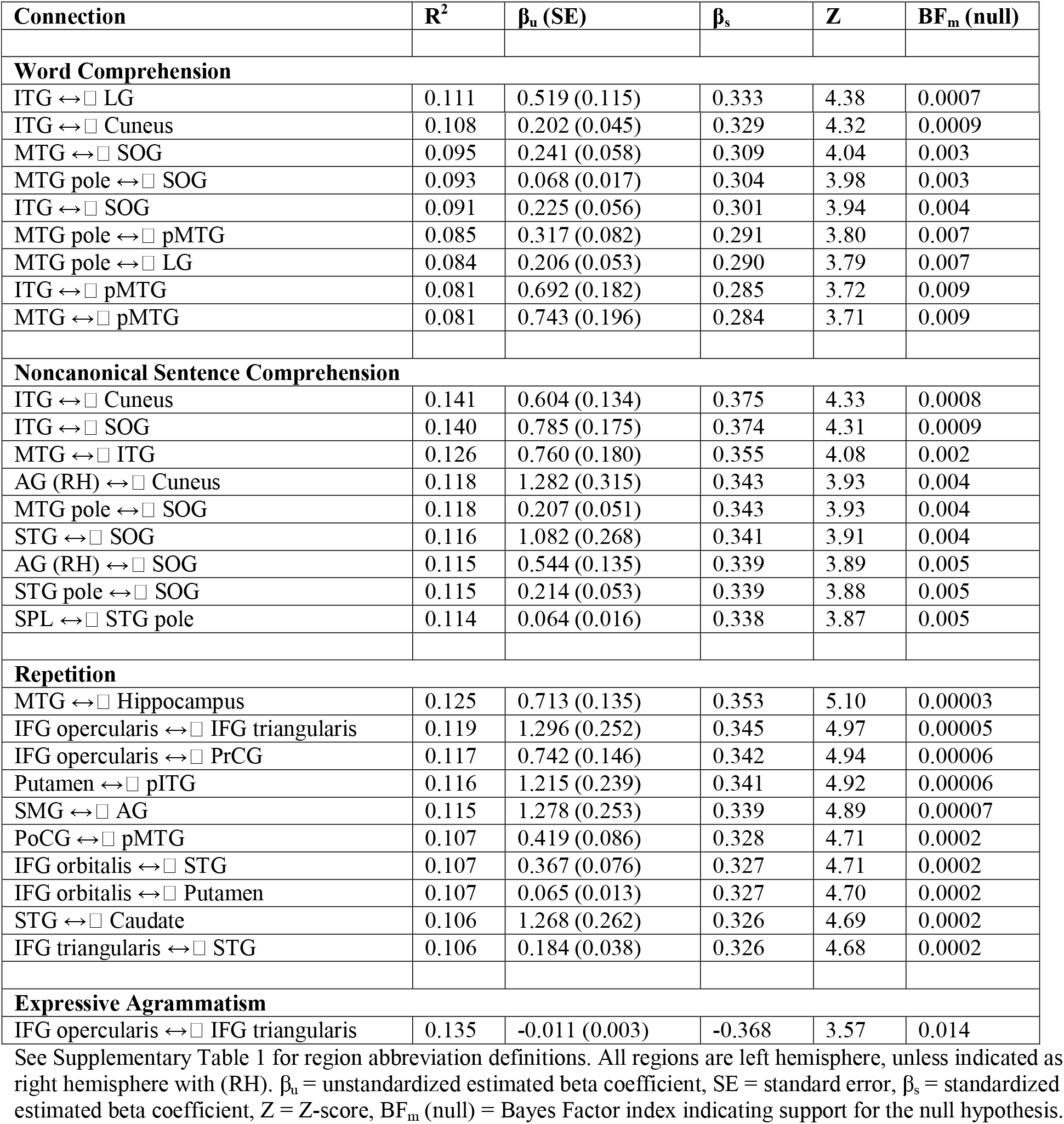
Top ten significant connections in the CLSM analyses.

| Connection | $R^2$ | $\beta_u$ (SE) | $\beta_s$ | Z | $BF_m$ (null) |
| --- | --- | --- | --- | --- | --- |
| **Word Comprehension** |  |  |  |  |  |
| ITG ↔ LG | 0.111 | 0.519 (0.115) | 0.333 | 4.38 | 0.0007 |
| ITG ↔ □ Cuneus | 0.108 | 0.202 (0.045) | 0.329 | 4.32 | 0.0009 |
| MTG ↔ □ SOG | 0.095 | 0.241 (0.058) | 0.309 | 4.04 | 0.003 |
| MTG pole ↔ □ SOG | 0.093 | 0.068 (0.017) | 0.304 | 3.98 | 0.003 |
| ITG ↔ □ SOG | 0.091 | 0.225 (0.056) | 0.301 | 3.94 | 0.004 |
| MTG pole ↔ □ pMTG | 0.085 | 0.317 (0.082) | 0.291 | 3.80 | 0.007 |
| MTG pole ↔ □ LG | 0.084 | 0.206 (0.053) | 0.290 | 3.79 | 0.007 |
| ITG ↔ □ pMTG | 0.081 | 0.692 (0.182) | 0.285 | 3.72 | 0.009 |
| MTG ↔ □ pMTG | 0.081 | 0.743 (0.196) | 0.284 | 3.71 | 0.009 |
| **Noncanonical Sentence Comprehension** |  |  |  |  |  |
| ITG ↔ □ Cuneus | 0.141 | 0.604 (0.134) | 0.375 | 4.33 | 0.0008 |
| ITG ↔ □ SOG | 0.140 | 0.785 (0.175) | 0.374 | 4.31 | 0.0009 |
| MTG ↔ □ ITG | 0.126 | 0.760 (0.180) | 0.355 | 4.08 | 0.002 |
| AG (RH) ↔ □ Cuneus | 0.118 | 1.282 (0.315) | 0.343 | 3.93 | 0.004 |
| MTG pole ↔ □ SOG | 0.118 | 0.207 (0.051) | 0.343 | 3.93 | 0.004 |
| STG ↔ □ SOG | 0.116 | 1.082 (0.268) | 0.341 | 3.91 | 0.004 |
| AG (RH) ↔ □ SOG | 0.115 | 0.544 (0.135) | 0.339 | 3.89 | 0.005 |
| STG pole ↔ □ SOG | 0.115 | 0.214 (0.053) | 0.339 | 3.88 | 0.005 |
| SPL ↔ □ STG pole | 0.114 | 0.064 (0.016) | 0.338 | 3.87 | 0.005 |
| **Repetition** |  |  |  |  |  |
| MTG ↔ □ Hippocampus | 0.125 | 0.713 (0.135) | 0.353 | 5.10 | 0.00003 |
| IFG opercularis ↔ □ IFG triangularis | 0.119 | 1.296 (0.252) | 0.345 | 4.97 | 0.00005 |
| IFG opercularis ↔ □ PrCG | 0.117 | 0.742 (0.146) | 0.342 | 4.94 | 0.00006 |
| Putamen ↔ □ pITG | 0.116 | 1.215 (0.239) | 0.341 | 4.92 | 0.00006 |
| SMG ↔ □ AG | 0.115 | 1.278 (0.253) | 0.339 | 4.89 | 0.00007 |
| PoCG ↔ □ pMTG | 0.107 | 0.419 (0.086) | 0.328 | 4.71 | 0.0002 |
| IFG orbitalis ↔ □ STG | 0.107 | 0.367 (0.076) | 0.327 | 4.71 | 0.0002 |
| IFG orbitalis ↔ □ Putamen | 0.107 | 0.065 (0.013) | 0.327 | 4.70 | 0.0002 |
| STG ↔ □ Caudate | 0.106 | 1.268 (0.262) | 0.326 | 4.69 | 0.0002 |
| IFG triangularis ↔ □ STG | 0.106 | 0.184 (0.038) | 0.326 | 4.68 | 0.0002 |
| **Expressive Agrammatism** |  |  |  |  |  |
| IFG opercularis ↔ □ IFG triangularis | 0.135 | -0.011 (0.003) | -0.368 | 3.57 | 0.014 |

By contrast, impaired Repetition was associated with a very large number of significant disconnections, within the frontal and temporal lobes as well as between frontal lobe and all other lobes, and between temporal lobe and all other lobes (there were no significant disconnections between parietal and occipital lobes). Expressive Agrammatism was associated with one significant disconnection in the frontal lobe, between IFG pars triangularis and IFG pars opercularis. These results underscore that the lack of significant frontal-based disconnections for Noncanonical Sentence Comprehension was not due to lack of statistical power or other methodological issues.

### 3.4 LSM and CLSM combined

Figure 4 and Supplementary Table 1 show the results of the combined analyses. The lesion-deficit correlations for all four middle and posterior temporal lobe regions (STG, MTG, pSTG, pMTG) for Word Comprehension remained statistically robust even when including the strongest relevant disconnections involving a more anterior temporal lobe region, MTG ↔□ SOG for the pSTG and pMTG and MTG pole SOG for the STG and MTG. Likewise, the lesion-deficit correlations for all the same middle and posterior temporal lobe regions (STG, MTG, pSTG, pMTG) for Noncanonical Sentence Comprehension remained statistically robust even when including the strongest relevant frontal disconnection (IFG orbitalis ↔□ Cuneus). Thus, there remains a strong independent association between damage to middle-posterior temporal lobe regions and comprehension deficits even when accounting for the strongest disconnections of ATL and the frontal lobe.

**Figure 4.**
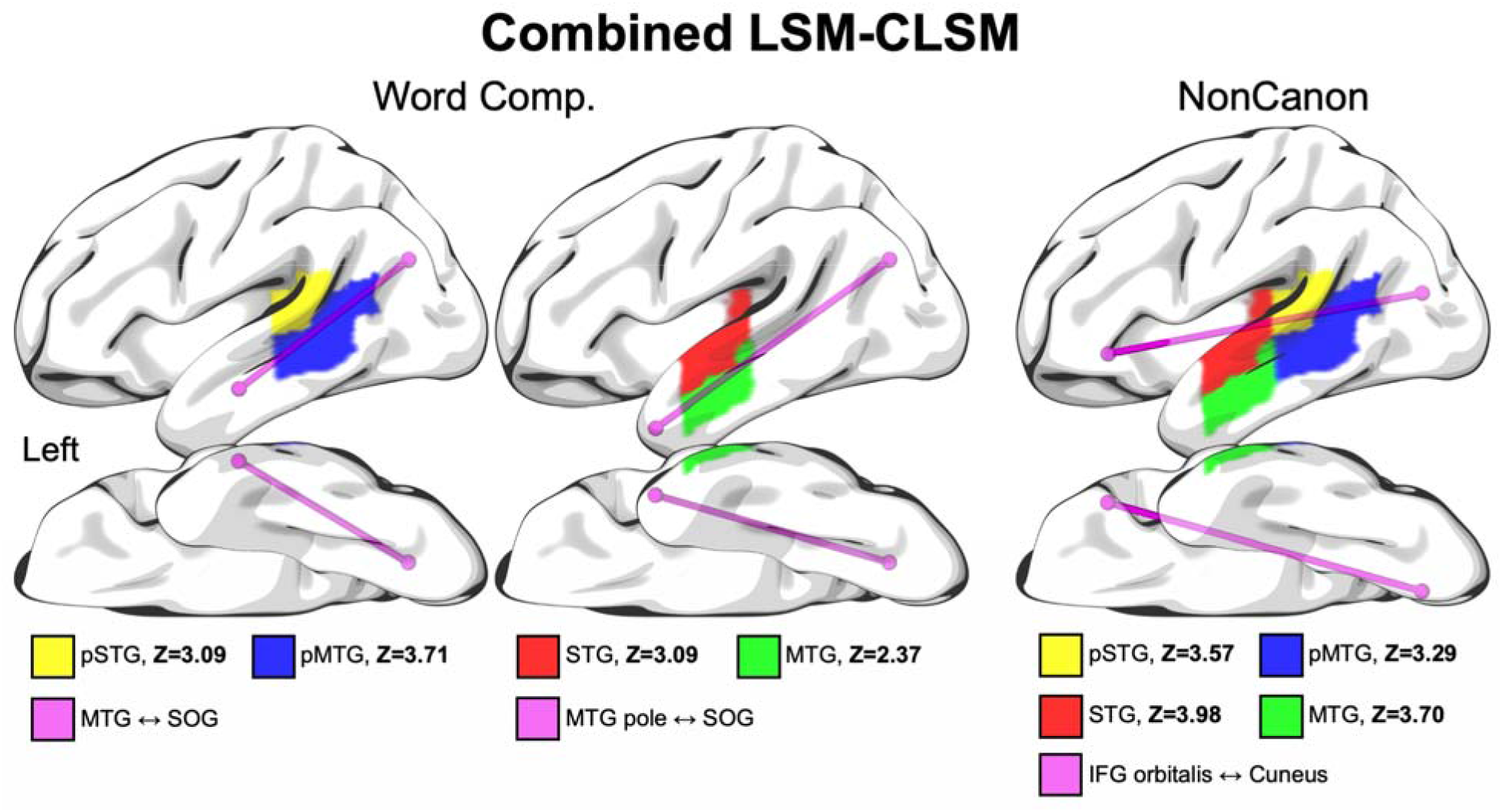
Combined LSM-CLSM analyses. Selected regions and connections for the analyses combining LSM and CLSM data, in which connection strength was included as a covariate for the linear regression analysis relating proportion damage within each region to the behavioral variable, including total lesion volume as a covariate. Z-scores for the association of damage to each region with comprehension deficits, after taking into account variance associated with connection strength, are shown at the bottom. All Z-scores are above 2.33, which corresponds to p < 0.01. Comp. = Comprehension, NonCanon = Noncanonical Sentence Comprehension, Z = Z-score, STG = superior temporal gyrus, MTG = middle temporal gyrus, pSTG = posterior superior temporal gyrus, pMTG = posterior middle temporal gyrus.

## 4 Discussion

Consistent with previous research, the lesion correlates of Word Comprehension (WAB-R auditory word recognition with PPT as a covariate) and Noncanonical Sentence Comprehension deficits in chronic post-stroke aphasia converged on the middle-posterior temporal lobe in our lesion symptom mapping (LSM) analyses, overlapping in the middle STG and middle-posterior STS. Because we (and Mesulam et al.^1^) used the PPT as a covariate in our analyses of Word Comprehension, variance associated with conceptual-semantic processing was likely greatly reduced, and thus this measure should highlight mechanisms up to and including lexical access. Given previous functional neuroimaging studies, we suggest that the middle STG is associated with phonetic and/or phonological mechanisms^60,61^ and the middle-posterior STS with lexical access^17^. The supplementary analyses of these measures including Repetition as a covariate resulted in similar lesion maps but with greatly reduced statistical power, without significant overlap but in nearly identical locations. The overlap of word and sentence comprehension deficits observed in this study is roughly consistent with the historical definition of Wernicke’s area, although it should be noted that robust overlap occurred more anteriorly than as often depicted (although see Wernicke^62^ for localization remarkably similar to what we observed here and in functional neuroimaging studies of lexical and syntactic processing). In general, we agree with others who have argued^3,63^ that the term “Wernicke’s area” should be avoided because of potential confusion about its localization stemming from the scattered historical anatomical localization of Wernicke’s area that most often focuses on more posterior temporal-parietal regions^1,2^. However, our results do reinforce the mostly uncontroversial conclusion that the temporal lobe regions associated with both word and sentence comprehension deficits in stroke-based aphasia are clearly posterior to the anterior locus of word comprehension deficits identified in the papers reported by Mesulam et al.^1,12^.

One of the main claims of Mesulam et al.^1^ is that damage to the territory associated with Wernicke’s area involves “double disconnection” in Wernicke’s aphasia. That is, they posited that word comprehension deficits follow from disconnection of the anterior temporal lobe (ATL), and sentence comprehension deficits follow from disconnection of frontal cortex, to various other regions of the brain, potentially including the right hemisphere. However, while our connectome-based lesion-symptom mapping (CLSM) analyses found that there were significant disconnections involving the temporal pole for both word and noncanonical sentence comprehension deficits, no significant frontal disconnections were associated with noncanonical sentence comprehension deficits. Our combined LSM and CLSM analyses further showed that, using the strongest sub-threshold frontal-based connection as a covariate, there were still robust effects of damage to middle and posterior temporal lobe regions. Thus, the fundamental prediction of Mesulam et al. that sentence comprehension deficits associated with damage to Wernicke’s area are explained by a frontal disconnection pattern was disconfirmed. Note that our results are unlikely to be due to lack of statistical power, as we identified significant frontal-temporal disconnections associated with speech repetition deficits, and a significant LSM effect of expressive agrammatism in frontal areas (particularly pMFG) as well as somewhat weaker effects in the ATL, with a significantly damaged connection between IFG pars opercularis and IFG pars triangularis.

The Mesulam et al. frontal-temporal disconnection proposal regarding sentence comprehension deficits has logical force if a primary cortical locus of syntactic comprehension were in the frontal lobe, as suggested by many authors^64–68^. Accordingly, we did find that agrammatic production deficits are associated with damage to the frontal lobe, including Broca’s area/posterior IFG, consistent with previous work^32,44,45,69–73^. In addition, functional neuroimaging studies of syntactic processing do reliably identify frontal activations including Broca’s area as indicated by meta-analyses of these studies^74,75^. However, recent theories propose that the systems for building hierarchical syntactic structure in *comprehension* are in the posterior temporal lobe, and not the frontal lobe^16,65^. The lack of a significant association between noncanonical sentence comprehension deficits and damage to the inferior or middle frontal lobe, and the lack of an association with significant frontal-temporal disconnections, provides evidence against the view that syntactic computations associated with sentence comprehension are processed in or around Broca’s area. Matchin and Hickok (2020) proposed that hierarchical syntax and lexical access are both subserved by the posterior temporal lobe, intimately intertwined, which is consistent with modern approaches to syntax positing “lexicalized” syntactic representations. Under this view, coincidence of the lesion correlates of lexical and syntactic processing in the posterior temporal lobe is expected. However, we must also keep in mind that the lack of frontal effects for syntactic comprehension might also be due to functional reorganization in chronic aphasia, and there is some evidence that people with chronic aphasia show enhanced activity for language processing in right IFG relative to healthy controls, as revealed by a recent meta-analysis^77^.

Importantly, our analyses of noncanonical sentence comprehension were matched to Mesulam et al.^1^ in assessing *overall* comprehension ability. Previous studies from our group in stroke-based aphasia that have examined *residual* performance on noncanonical structures after controlling for performance to canonical structures have associated deficits with damage to frontal regions^30,78^ (with other studies showing conflicting results^25,44,45^). Residual performance on noncanonical sentence comprehension after controlling for canonical sentence comprehension likely highlights executive function resources that are necessary for processing particularly complex structures and revising sentence interpretation^79–85^, thus some implication of frontal regions that are associated with these supporting mechanisms is expected^21,86–89^. The existence of sub-threshold frontal disconnections provides some support for this account.

One prediction of Mesulam et al.^1^ was partially confirmed: word comprehension deficits involved disconnection of anterior temporal regions. The disconnections we found between anterior temporal regions and other temporal lobe regions and visual cortex were not unique to word comprehension deficits, as similar disconnections were associated with noncanonical sentence comprehension deficits. However, we take the primary claim of Mesulam et al. to be not that disconnections of ATL are associated with word comprehension impairments, but rather that these disconnections *explain* the association of posterior temporal lobe damage with these impairments. Critically, our combined LSM-CLSM analyses revealed that damage to middle and posterior temporal lobe regions was robustly associated with word comprehension impairments even when including the strongest disconnections involving a more anterior temporal lobe region. In addition, we also tested the possibility raised by Mesulam et al. that word comprehension impairments are associated with disconnection of right hemisphere regions, but no significant right hemisphere disconnections were identified.

The Mesulam et al. hypothesis would also predict that damage to the ATL itself should be associated with word comprehension deficits. However, we note that the corrected voxel-based LSM analyses of Word Comprehension (when controlling for conceptual-semantic processing via the PPT covariate) did not reveal an association with ATL damage. This lack of a robust ATL effect for Word Comprehension is unlikely to be due to statistical power, as we had adequate lesion coverage in this area and did identify significant effects for Expressive Agrammatism there. Overall, these combined results speak against the disconnection hypothesis of Mesulam et al. They suggest that the ATL may play a supporting role for word comprehension by retrieving conceptual-semantic representations which are necessary to perform a picture task in some contexts, and thus ATL disconnection contributes in a minor way to the impaired word comprehension performance seen in many patients with post-stroke aphasia involving damage to the posterior temporal lobe.

Under Mesulam et al.’s hypothesis, Wernicke’s area (broadly construed) plays a key role in phonological short-term/working memory, which is in line with their reported association between repetition deficits and posterior temporal cortical degeneration^12^. Consistent with this, in our study, repetition deficits were associated with arcuate fasciculus and posterior temporal damage and multiple frontal-parietal-temporal disconnections. Furthermore, deficits in noncanonical sentence comprehension (but not auditory word comprehension), were associated with a significant disconnection between the parietal and temporal lobe. This converges with previous reports of associations between deficits in complex sentence comprehension and inferior parietal lobe damage in chronic stroke patients with aphasia^22,23,25,26,31,44,78,90^, as well as functional neuroimaging studies finding activation for sensory-motor integration in this vicinity^37,39,91^. It is possible that these disconnections reflect the additional phonological short term memory resources that are useful in parsing sentence structure that are not typically required for word comprehension^80–83,92–94,94^, which may explain why the significant correlation between word and sentence comprehension abilities was only moderate in strength.

Task difference is a major factor which helps explain the discrepancy between the lesion localization for word comprehension deficits we report here (and generally in the stroke aphasia literature) and those of Mesulam et al.^1,12^. The word comprehension task used by Mesulam et al. consists of moderately difficult items of the Peabody Picture Vocabulary Task (PPVT)^95^, which involve much more complex semantic inference than the task we used, the auditory word recognition subtest of the WAB-R^41^, likely resulting in considerable variance due to conceptual-semantic processing despite including the PPT as a covariate. Wernicke and Lichtheim thought of Wernicke’s area as the key node for “auditory word memories”^6,7,62^, which can be interpreted in modern linguistic terms as phonological and lexical processing, with conceptual-semantic representations widely distributed elsewhere in the brain. The WAB-R auditory word recognition test is therefore more appropriate to assess this idea, as it does not require fine-grained visual processing and object recognition, and limits semantic processing demands, unlike the PPVT. Future studies should examine the relationship between these word comprehension measures and the results obtained in lesion-symptom mapping analyses in both PPA and stroke-based aphasia.

In addition, PPA is a progressive neurodegenerative disease which has key differences from stroke based aphasia, evolving to ultimately affect a number of cognitive domains^96,97^. Functional neuroimaging studies of people with the semantic variant of PPA, associated primarily with ATL atrophy, have shown abnormal activation patterns and functional connectivity in regions outside the ATL^98–104^. This suggests that the neuropathology of semantic PPA is more widespread than what shows up in gross measures of grey matter atrophy and may affect a network (potentially including middle and posterior temporal cortex) that contributes to word comprehension impairments. Individuals with PPA show neural degeneration far beyond the areas of peak atrophy (as seen in the figures in Mesulam et al.^1^), as well as disrupted functional networks beyond the sites of major neural degeneration. Longitudinal MRI and word comprehension testing reveals that deterioration in auditory word comprehension over time in PPA is associated with within-individual atrophy in left middle temporal cortex, left angular gyrus, and right inferior and middle temporal cortex^105^. Moreover, the word level comprehension deficits seen in stroke patients and the semantic variant of PPA might have qualitative differences. Word comprehension deficits in stroke may be more commonly due to deficits in basic linguistic processing, i.e. at the phonological and lexical/lemma levels, whereas word comprehension deficits in PPA may be more commonly due to deficits in accessing finegrained semantic features of words downstream from lexical access. These may be more completely disrupted in semantic PPA due to the pattern of neurodegeneration than is seen with strokes impinging on this area, or in resections.

Under our account, in which the middle-posterior temporal lobe plays a dominant causal role in both lexical access and syntactic comprehension, one would expect a strong association between word and sentence comprehension deficits and degeneration of the middle and posterior temporal lobe in PPA. However, this is not what is seen, as patients with the logopenic variant of PPA and posterior temporal-parietal degeneration do not typically show notable word comprehension deficits^1,106–108^ (but see Bonner & Grossman^109^). Also, Mesulam et al.^1^ do not report a notable association of atrophy in middle-posterior temporal lobe with deficits in noncanonical sentence comprehension. Why not? Some of the present authors have previously suggested that logopenic PPA does not necessarily involve complete degeneration of posterior temporal-parietal cortex, and that remaining neurons may be sufficient to perform the basic phonological and lexical access functions^8^. Additionally, some models posit an important role for the right hemisphere in phonological processing and lexical access^110–112^, which could sufficiently compensate for left hemisphere degeneration. This is consistent with recent demonstrations of small but significant word comprehension deficits in right hemisphere stroke^113^.

One area of agreement between Mesulam et al. and our data is that damage or degeneration of the ATL is not implicated in noncanonical sentence comprehension deficits. However, we did find significant disconnections of the temporal pole ROIs related to impaired noncanonical sentence comprehension deficits. This is consistent with previous reports implicating damage to anterior temporal lobe in grammatical processing deficits^22,31^. Nevertheless, we suggest that the (relatively less common) implication of ATL damage or disconnection in sentence comprehension deficits likely reflects semantic rather than syntactic deficits^21,63^, which is consistent with functional neuroimaging data^114–118^.

Overall, our combined LSM and CLSM results speak against the hypothesis of Mesulam et al. (2015) of a double disconnection syndrome underlying word and sentence/syntactic comprehension deficits in post-stroke aphasia. Rather, our results support the classical concept of Wernicke’s area as directly supporting both word and sentence comprehension, although our results do suggest that anterior temporal lobe and inferior parietal networks bolster core linguistic processing through semantic and phonological working memory resources, respectively. The discrepancy between our results and those of Mesulam et al. from PPA might be explained by differences between language deficits in post-stroke aphasia versus PPA or differences in tests used to assess comprehension, or both. Alternatively, the conclusions based on PPA might be unfounded, because they were based on regions of peak atrophy associated with errors, rather than considering other areas of degeneration or dysfunction that might be responsible for deficits. Irrespective of the account of the discrepant results, our data provide strong evidence for the major role of the middle STG and middle-posterior STS in both word and sentence comprehension.

## Acknowledgments

We would also like to thank Alexandra Basilakos and Brielle C. Stark for assistant with rating of speech samples, Leigh Ann Spell, Allison Croxton, Anna Doyle, Michele Martin, Katie Murphy, and Sara Sayers for their assistance with data collection, and graduate student clinicians in the Aphasia Lab for transcribing and coding speech samples. Finally, we would like to thank two anonymous reviewers for their fruitful analysis suggestions.

## Supplementary Data

Supplementary tables and figures can be found in the supplementary data file.

