## Supplementary Data for "The Wernicke conundrum revisited: evidence from connectome-based lesion-symptom mapping"

**Supplementary Materials**

Figures 1 and 2 from the main body of the manuscript describe LSM analyses including our two primary measures of interest, Word Comprehension (WAB-R Auditory Word Recognition with Pyramids and Palm Trees, PPT, as a covariate) and Noncanonical Sentence Comprehension. Here we report supplementary analyses of these measures including an additional covariate, Repetition (WAB-R Repetition score), in an attempt to control for phonological processing and focus on lexical and syntactic levels of representation. The results are shown in Supplementary Figure 1.

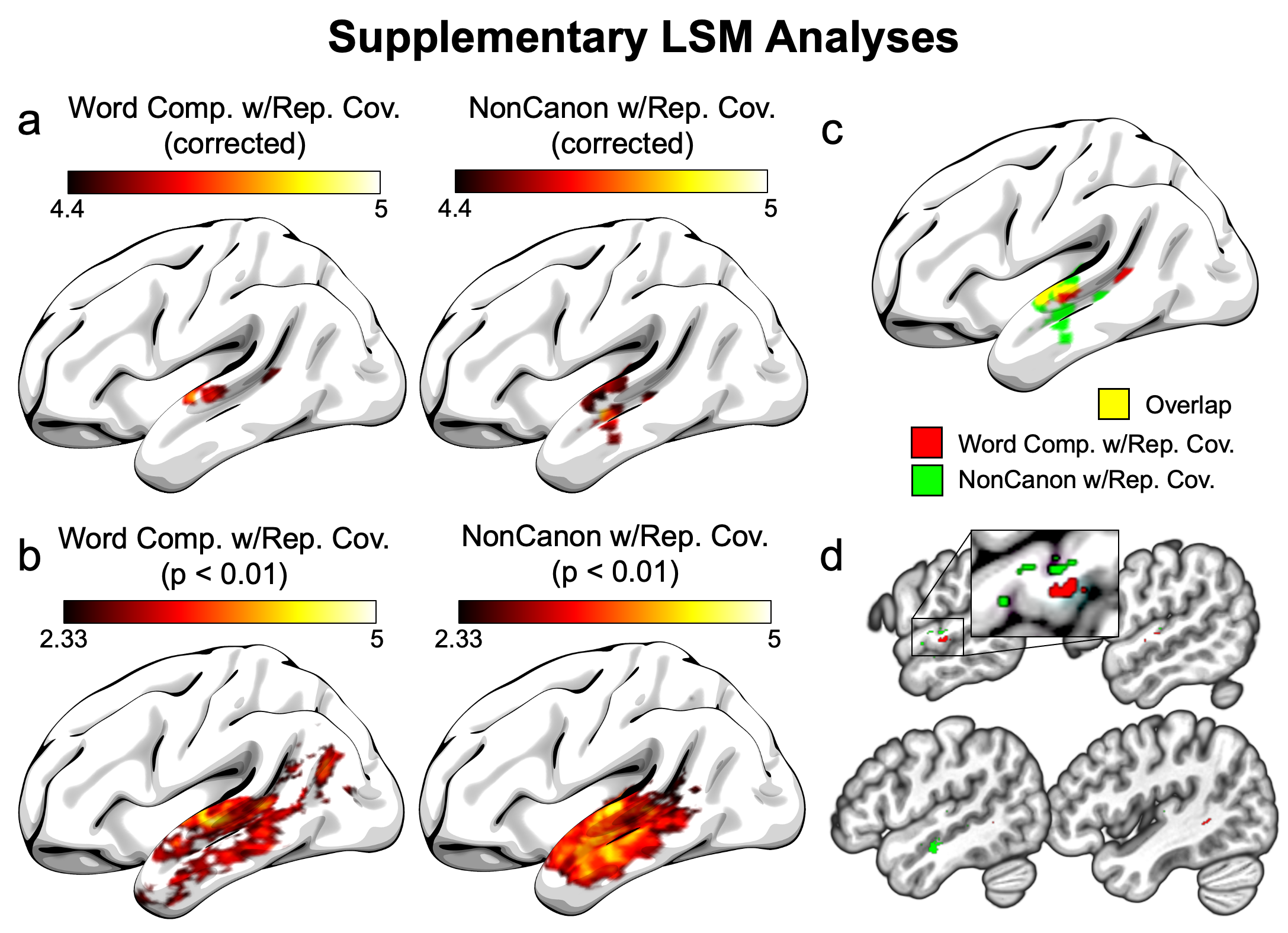

Supplementary Figure 1. Supplementary voxel-based lesion-symptom mapping (LSM) results for Word Comprehension (WAB-R Auditory Word Recognition with the PPT covariate) and Noncanonical Sentence Comprehension incorporating Repetition scores as a covariate. a) Analyses with a permutation correction for multiple comparisons (10,000 permutations, corrected p < 0.05), with results spatially smoothed for improved visibility. b) Unsmoothed analyses using an uncorrected voxel-wise threshold of p < 0.01 (one-tailed). c) Smoothed surface rendering of the overlap (yellow) of the corrected lesion maps for Word Comprehension (red) and Noncanonical Sentence Comprehension (green). d) b) Unsmoothed slices of the volumetric (non-surface-rendered) overlap. Inset shows a zoomed-in view of the portion of the slice surrounded by the rectangle.

No frontal-temporal disconnections survived the CLSM analyses when corrected for multiple comparisons. Selected frontal-temporal connections of interest, involving all three sub-regions of the IFG as well as the pMTG region that was significantly associated with Expressive Agrammatism and the four middle and posterior temporal lobe regions (STG, MTG, pSTG, pMTG) associated with Noncanonical sentence comprehension are shown in Supplementary Table 1. We subsequently selected four of these sub-threshold connections for use in the combined LSM-CLSM analyses.

**Supplementary Table 1 Sub-threshold frontal-temporal connections for Noncanonical sentence comprehension.**

| **Connection** | **R^2^** | **β_u_ (SE)** | **β_s_** | **Z** | **BF_m_ (null)** |
| --- | --- | --- | --- | --- | --- |
| pMFG ↔︎ STG | 0.022 | 0.019 (0.011) | 0.147 | 1.65 | 1.518 |
| pMFG ↔︎ MTG | 0.019 | 0.042 (0.027) | 0.139 | 1.55 | 1.746 |
| pMFG ↔︎ pSTG | 0.009 | 0.051 (0.047) | 0.096 | 1.07 | 3.114 |
| pMFG ↔︎ pMTG | 0.005 | 0.231 (0.298) | 0.069 | 0.77 | 4.009 |
| **IFG opercularis** ↔︎ **STG** | **0.055** | **0.084 (0.031)** | **0.236** | **2.66** | **0.205** |
| IFG opercularis ↔︎ MTG | 0.030 | 0.231 (0.119) | 0.172 | 1.93 | 0.954 |
| IFG opercularis ↔︎ pSTG | 0.008 | 0.189 (0.185) | 0.091 | 1.02 | 3.283 |
| IFG opercularis ↔︎ pMTG | 0.009 | 0.405 (0.385) | 0.094 | 1.05 | 3.186 |
| IFG orbitalis ↔︎ STG | 0.010 | 0.128 (0.115) | 0.100 | 1.11 | 2.997 |
| IFG orbitalis ↔︎ MTG | 0.029 | 0.142 (0.073) | 0.172 | 1.92 | 0.965 |
| **IFG orbitalis** ↔︎ **pSTG** | **0.017** | **0.080 (0.054)** | **0.132** | **1.47** | **1.949** |
| **IFG orbitalis** ↔︎ **pMTG** | **0.020** | **0.227 (0.142)** | **0.143** | **1.59** | **1.648** |
| IFG triangularis ↔︎ STG | 0.046 | 0.142 (0.058) | 0.215 | 2.42 | 0.355 |
| **IFG triangularis** ↔︎ **MTG** | **0.042** | **0.237 (0.101)** | **0.205** | **2.31** | **0.456** |
| IFG triangularis ↔︎ pSTG | 0.016 | 0.201 (0.142) | 0.126 | 1.40 | 2.138 |
| IFG triangularis ↔︎ pMTG | 0.013 | 0.359 (0.276) | 0.116 | 1.29 | 2.455 |

The four strongest connections, each involving one of the four temporal lobe regions significantly identified in the LSM analysis of Noncanonical sentence comprehension, selected for combined LSM-CLSM analyses are shown in bold. See Figure 3 for region abbreviation definitions. All regions are left hemisphere. β_u_ = unstandardized estimated beta coefficient, SE = standard error, β_s_ = standardized estimated beta coefficient, Z = Z-score, BF_m_ (null) = Bayes Factor index indicating support for the null hypothesis.

The statistical details of the combined LSM-CLSM analyses detailed in section 2.3.4, with the results summarized in section 3.4, are provided in Supplementary Table 2. The analysis of Word Comprehension (WAB-R Auditory Word Recognition with PPT covariate) assessed the contribution of lesion volume in the two posterior temporal lobe regions identified in our ROI-based LSM analyses, controlling for the strongest connection involving that region to a more anterior temporal lobe region identified in the CLSM analyses as well as lesion volume. The analysis of Noncanonical (Noncanonical sentence comprehension) assessed the contribution of lesion volume in all of the four middle and posterior temporal lobe regions identified in our ROI-based LSM analyses, controlling for the strongest sub-threshold connection involving that region identified in the CLSM analyses as well as lesion volume.

**Supplementary Table 2 LSM and CLSM combined**

| **Analysis** | **R^2^** | **β_u_ (SE)** | **β_s_** | **Z** | **BF_m_ (null)** |
| --- | --- | --- | --- | --- | --- |
| **Word Comprehension** | | | | | |
| pSTG, controlling for STG ↔︎ pSTG | 0.203 | -8.568 (2.471) | -0.369 | 3.21 | 1.274e-6 |
| pMTG, controlling for MTG pole ↔︎ pMTG | 0.256 | -9.103 (2.139) | -0.362 | 3.98 | 2.719e-8 |
| **Noncanonical** | | | | | |
| STG, controlling for IFG opercularis ↔︎ STG | 0.254 | -4.736 (1.017) | -0.507 | 4.31 | 2.750e-6 |
| MTG, controlling for IFG triangularis ↔︎ MTG | 0.232 | -4.474 (1.047) | -0.414 | 3.96 | 1.691e-5 |
| pSTG, controlling for IFG orbitalis ↔︎ pSTG | 0.205 | -3.693 (0.927) | -0.427 | 3.68 | 5.599e-5 |
| pMTG, controlling for IFG orbitalis ↔︎ pMTG | 0.196 | -3.537 (0.931) | -0.375 | 3.51 | 1.493e-4 |

See Figure 3 for region abbreviation definitions. All regions are left hemisphere. β_u_ = unstandardized estimated beta coefficient, SE = standard error, β_s_ = standardized estimated beta coefficient, Z = Z-score, BF_m_ (null) = Bayes Factor index indicating support for the null hypothesis.

In order to illustrate the goodness of linear fits to our ROI-based LSM and CLSM data, Supplementary Figures 2 and 3 report scatterplots of the individual subject data used in these analyses.

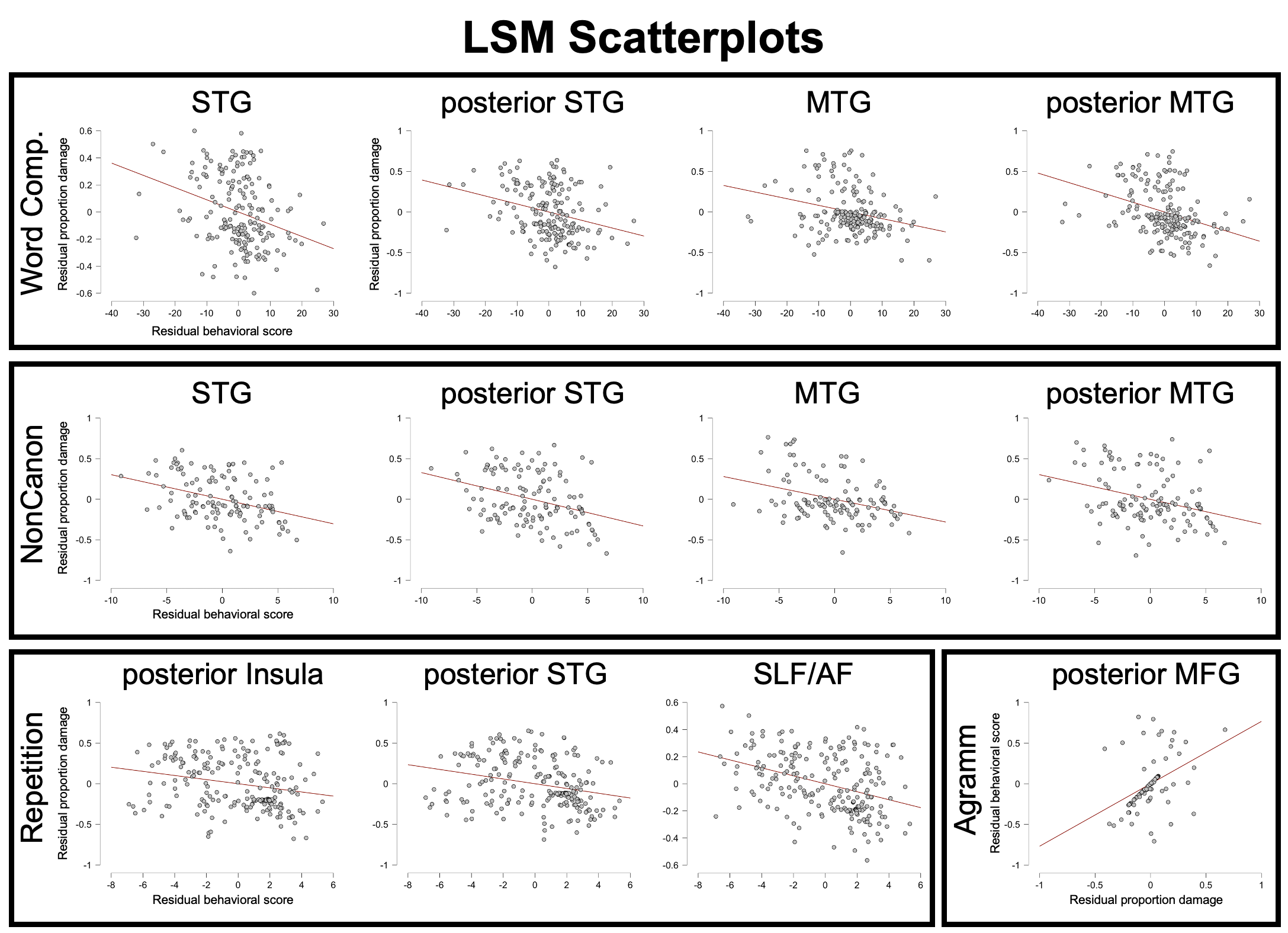

Supplementary Figure 2. Scatterplots of the ROI-based LSM analyses for each behavioral measure, reflecting residual values after the effect of lesion volume is regressed out. See Figure 3 for region abbreviation definitions. Comp = Comprehension; NonCanon = Noncanonical Sentence Comprehension; Agramm = Expressive Agrammatism.

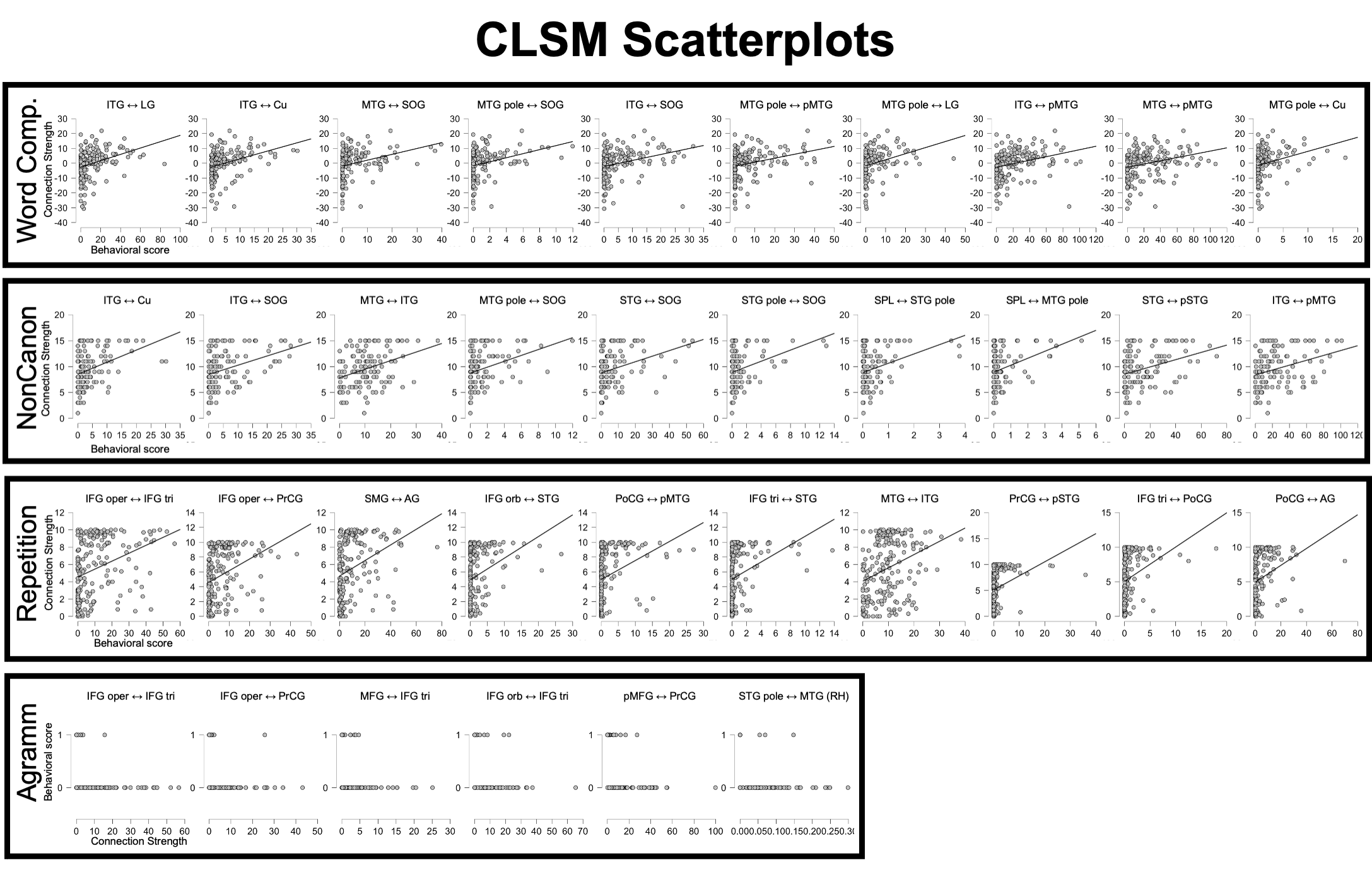

Supplementary Figure 3. Scatterplots of the top ten significant disconnections from the CLSM analyses for each behavioral measure. Comp = Comprehension; NonCanon = Noncanonical Sentence Comprehension; Agramm = Expressive Agrammatism. See Figure 3 for region abbreviation definitions.

In Table 4, we reported the top ten significant connections between regions from the JHU atlas (Faria et al., 2012) to which damage was associated with each of the four primary behavioral measures in our connectome lesion-symptom mapping (CLSM) analyses. Supplementary Table 3 lists the entire set of significant connections, as well as the significant connections for the two supplemental behavioral measures, WAB-R Word Comprehension with PPT and WAB-R Repetition Covariates and Noncanonical Sentence Comprehension with WAB-R Repetition Covariate.

**Supplementary Table 3. Full list of significant connections in CLSM analyses.**

| Connection | R^2^ | β_u_ (SE) | β_s_ | Z | BF_m_ (null) |
| --- | --- | --- | --- | --- | --- |
| **Word Comprehension** | | | | | |
| ITG ↔︎ LG | 0.111 | 0.519 (0.115) | 0.333 | 4.38 | 0.0007 |
| ITG ↔︎ Cu | 0.108 | 0.202 (0.045) | 0.329 | 4.32 | 0.0009 |
| MTG ↔︎ SOG | 0.095 | 0.241 (0.058) | 0.309 | 4.04 | 0.003 |
| MTG pole ↔︎ SOG | 0.093 | 0.068 (0.017) | 0.304 | 3.98 | 0.003 |
| ITG ↔︎ SOG | 0.091 | 0.225 (0.056) | 0.301 | 3.94 | 0.004 |
| MTG pole ↔︎ pMTG | 0.085 | 0.317 (0.082) | 0.291 | 3.80 | 0.007 |
| MTG pole ↔︎ LG | 0.084 | 0.206 (0.053) | 0.290 | 3.79 | 0.007 |
| ITG ↔︎ pMTG | 0.081 | 0.692 (0.182) | 0.285 | 3.72 | 0.009 |
| MTG ↔︎ pMTG | 0.081 | 0.743 (0.196) | 0.284 | 3.71 | 0.009 |
| MTG pole ↔︎ Cu | 0.073 | 0.077 (0.022) | 0.271 | 3.53 | 0.017 |
| STG ↔︎ SOG | 0.073 | 0.294 (0.082) | 0.270 | 3.51 | 0.018 |
| MTG pole ↔︎ IOG | 0.065 | 0.217 (0.064) | 0.255 | 3.31 | 0.034 |
| STG ↔︎ Cu | 0.065 | 0.176 (0.052) | 0.254 | 3.30 | 0.035 |
| STG ↔︎ pSTG | 0.065 | 0.400 (0.119) | 0.254 | 3.30 | 0.035 |
| STG ↔︎ pMTG | 0.063 | 0.321 (0.097) | 0.250 | 3.25 | 0.042 |
| MTG pole ↔︎ MOG | 0.062 | 0.148 (0.045) | 0.249 | 3.23 | 0.044 |
| SOG ↔︎ pMTG | 0.061 | 0.258 (0.079) | 0.248 | 3.21 | 0.047 |
| **Noncanonical Sentence Comprehension** | | | | | |
| ITG ↔︎ Cu | 0.141 | 0.604 (0.134) | 0.375 | 4.33 | 0.0008 |
| ITG ↔︎ SOG | 0.140 | 0.785 (0.175) | 0.374 | 4.31 | 0.0009 |
| MTG ↔︎ ITG | 0.126 | 0.760 (0.180) | 0.355 | 4.08 | 0.002 |
| MTG pole ↔︎ SOG | 0.118 | 0.207 (0.051) | 0.343 | 3.93 | 0.004 |
| STG ↔︎ SOG | 0.116 | 1.082 (0.268) | 0.341 | 3.91 | 0.004 |
| STG pole ↔︎ SOG | 0.115 | 0.214 (0.053) | 0.339 | 3.88 | 0.005 |
| SPL ↔︎ STG pole | 0.114 | 0.064 (0.016) | 0.338 | 3.87 | 0.005 |
| SPL ↔︎ MTG pole | 0.112 | 0.083 (0.021) | 0.334 | 3.82 | 0.006 |
| STG ↔︎ pSTG | 0.108 | 1.533 (0.396) | 0.328 | 3.75 | 0.008 |
| ITG ↔︎ pMTG | 0.106 | 2.262 (0.590) | 0.326 | 3.72 | 0.008 |
| IFG orbitalis ↔︎ Cu | 0.103 | 0.096 (0.025) | 0.321 | 3.66 | 0.010 |
| STG pole ↔︎ Cu | 0.103 | 0.183 (0.049) | 0.321 | 3.66 | 0.010 |
| MTG ↔︎ pMTG | 0.100 | 2.317 (0.624) | 0.316 | 3.61 | 0.013 |
| ITG ↔︎ MOG | 0.099 | 1.795 (0.485) | 0.315 | 3.60 | 0.013 |
| PoCG ↔︎ STG pole | 0.099 | 0.023 (0.006) | 0.315 | 3.59 | 0.013 |
| LFOG ↔︎ Cu | 0.093 | 0.057 (0.016) | 0.306 | 3.48 | 0.019 |
| PoCG ↔︎ MTG_pole | 0.091 | 0.018 (0.005) | 0.301 | 3.43 | 0.023 |
| SMG ↔︎ MTG pole | 0.090 | 0.043 (0.012) | 0.299 | 3.41 | 0.024 |
| ITG ↔︎ LG | 0.089 | 1.242 (0.357) | 0.298 | 3.39 | 0.025 |
| MTG pole ↔︎ Cu | 0.089 | 0.204 (0.059) | 0.298 | 3.39 | 0.026 |
| ITG ↔︎ FuG | 0.084 | 1.687 (0.501) | 0.289 | 3.28 | 0.036 |
| **WAB-R repetition** | | | | | |
| IFG opercularis ↔︎ IFG triangularis | 0.119 | 1.296 (0.252) | 0.345 | 4.97 | 0.00005 |
| IFG opercularis ↔︎ PrCG | 0.117 | 0.742 (0.146) | 0.342 | 4.94 | 0.00006 |
| SMG ↔︎ AG | 0.115 | 1.278 (0.253) | 0.339 | 4.89 | 0.00007 |
| IFG orbitalis ↔︎ STG | 0.107 | 0.367 (0.076) | 0.327 | 4.71 | 0.0002 |
| PoCG ↔︎ pMTG | 0.107 | 0.419 (0.086) | 0.328 | 4.71 | 0.0002 |
| IFG triangularis ↔︎ STG | 0.106 | 0.184 (0.038) | 0.326 | 4.68 | 0.0002 |
| MTG ↔︎ ITG | 0.102 | 0.662 (0.141) | 0.319 | 4.58 | 0.0003 |
| PrCG ↔︎ pSTG | 0.098 | 0.363 (0.078) | 0.314 | 4.50 | 0.0004 |
| IFG triangularis ↔︎ PoCG | 0.095 | 0.192 (0.042) | 0.309 | 4.42 | 0.0006 |
| PoCG ↔︎ AG | 0.092 | 0.758 (0.170) | 0.304 | 4.35 | 0.0008 |
| IFG triangularis ↔︎ SOG | 0.091 | 0.072 (0.016) | 0.302 | 4.32 | 0.0009 |
| IFG triangularis ↔︎ Cu | 0.090 | 0.037 (0.008) | 0.301 | 4.30 | 0.001 |
| IFG triangularis ↔︎ LG | 0.088 | 0.071 (0.016) | 0.297 | 4.25 | 0.001 |
| IFG orbitalis ↔︎ LG | 0.085 | 0.216 (0.050) | 0.292 | 4.18 | 0.002 |
| IFG opercularis ↔︎ PoCG | 0.085 | 0.222 (0.052) | 0.291 | 4.17 | 0.002 |
| STG ↔︎ SOG | 0.084 | 0.808 (0.191) | 0.289 | 4.13 | 0.002 |
| IFG triangularis ↔︎ PrCG | 0.083 | 0.372 (0.088) | 0.288 | 4.11 | 0.002 |
| IFG orbitalis ↔︎ MTG | 0.082 | 0.208 (0.050) | 0.285 | 4.08 | 0.002 |
| IFG orbitalis ↔︎ FuG | 0.080 | 0.123 (0.030) | 0.283 | 4.04 | 0.003 |
| PrCG ↔︎ SMG | 0.079 | 0.796 (0.195) | 0.280 | 4.00 | 0.003 |
| SPL ↔︎ pMTG | 0.078 | 0.764 (0.188) | 0.279 | 3.98 | 0.004 |
| IFG orbitalis ↔︎ SOG | 0.078 | 0.118 (0.029) | 0.279 | 3.97 | 0.004 |
| PoCG ↔︎ SMG | 0.077 | 1.108 (0.274) | 0.277 | 3.95 | 0.004 |
| SPL ↔︎ pSTG | 0.077 | 0.898 (0.223) | 0.277 | 3.95 | 0.004 |
| PrCG ↔︎ pMTG | 0.076 | 0.861 (0.214) | 0.276 | 3.94 | 0.004 |
| SPL ↔︎ STG pole | 0.076 | 0.046 (0.011) | 0.276 | 3.93 | 0.004 |
| STG (RH) ↔︎ STG pole | 0.076 | 0.011 (0.003) | 0.276 | 3.93 | 0.004 |
| STG ↔︎ MTG | 0.074 | 0.567 (0.143) | 0.272 | 3.88 | 0.005 |
| PoCG ↔︎ STG pole | 0.074 | 0.017 (0.004) | 0.271 | 3.87 | 0.005 |
| pMFG ↔︎ IFG opercularis | 0.073 | 1.138 (0.289) | 0.270 | 3.85 | 0.006 |
| ITG ↔︎ LG | 0.072 | 1.114 (0.287) | 0.267 | 3.81 | 0.007 |
| pMFG ↔︎ PrCG | 0.071 | 1.178 (0.305) | 0.266 | 3.78 | 0.007 |
| IFG triangularis ↔︎ SMG | 0.070 | 0.280 (0.073) | 0.264 | 3.76 | 0.008 |
| RG ↔︎ SOG | 0.068 | 0.031 (0.008) | 0.261 | 3.71 | 0.009 |
| SMG ↔︎ pMTG | 0.066 | 0.727 (0.195) | 0.257 | 3.66 | 0.011 |
| IFG opercularis ↔︎ Cu | 0.064 | 0.009 (0.002) | 0.253 | 3.60 | 0.014 |
| AG ↔︎ pMTG | 0.064 | 1.819 (0.497) | 0.253 | 3.60 | 0.014 |
| SPL ↔︎ AG | 0.064 | 1.874 (0.514) | 0.252 | 3.58 | 0.015 |
| SPL ↔︎ MTG | 0.063 | 0.229 (0.063) | 0.252 | 3.58 | 0.015 |
| MFG ↔︎ STG | 0.062 | 0.056 (0.016) | 0.249 | 3.54 | 0.017 |
| IFG opercularis ↔︎ LG | 0.061 | 0.016 (0.004) | 0.247 | 3.51 | 0.019 |
| SPL ↔︎ SMG | 0.060 | 1.413 (0.398) | 0.246 | 3.49 | 0.020 |
| PoCG ↔︎ pSTG | 0.060 | 0.153 (0.043) | 0.245 | 3.48 | 0.021 |
| PrCG ↔︎ AG | 0.060 | 1.109 (0.314) | 0.245 | 3.47 | 0.021 |
| IFG triangularis ↔︎ FuG | 0.059 | 0.048 (0.014) | 0.243 | 3.45 | 0.023 |
| STG pole ↔︎ MTG (RH) | 0.059 | 0.008 (0.002) | 0.242 | 3.44 | 0.024 |
| IFG opercularis ↔︎ SMG | 0.058 | 0.375 (0.108) | 0.240 | 3.41 | 0.027 |
| IFG opercularis ↔︎ pSTG | 0.057 | 0.415 (0.120) | 0.239 | 3.40 | 0.028 |
| IFG triangularis ↔︎ pSTG | 0.057 | 0.321 (0.094) | 0.238 | 3.37 | 0.030 |
| MFG ↔︎ MTG | 0.055 | 0.039 (0.012) | 0.234 | 3.32 | 0.035 |
| SPL ↔︎ STG | 0.054 | 0.550 (0.165) | 0.232 | 3.29 | 0.038 |
| STG pole ↔︎ pSTG (RH) | 0.053 | 0.020 (0.006) | 0.231 | 3.28 | 0.040 |
| IFG orbitalis ↔︎ PoCG | 0.052 | 0.055 (0.017) | 0.228 | 3.23 | 0.046 |
| IFG orbitalis ↔︎ Cu | 0.052 | 0.059 (0.018) | 0.228 | 3.23 | 0.046 |
| ITG ↔︎ SOG | 0.052 | 0.439 (0.134) | 0.228 | 3.22 | 0.047 |
| **Expressive Agrammatism** | | | | | |
| IFG opercularis ↔︎ IFG triangularis | 0.135 | -0.011 (0.003) | -0.368 | 3.57 | 0.014 |
| IFG opercularis ↔︎ PrCG | 0.091 | -0.014 (0.005) | -0.302 | 2.89 | 0.105 |
| MFG ↔︎ IFG triangularis | 0.086 | -0.024 (0.008) | -0.294 | 2.81 | 0.128 |
| IFG orbitalis ↔︎ IFG triangularis | 0.080 | -0.010 (0.004) | -0.284 | 2.71 | 0.166 |
| pMFG ↔︎ PrCG | 0.080 | -0.007 (0.002) | -0.282 | 2.69 | 0.172 |
| STG pole ↔︎ MTG (RH) | 0.074 | -1.574 (0.594) | -0.272 | 2.59 | 0.221 |
| **Word Comprehension with Repetition covariate** | | | | | |
| MTG pole ↔︎ pMTG | 0.092 | 0.356 (0.087) | 0.303 | 3.97 | 0.004 |
| MTG pole ↔︎ SOG | 0.074 | 0.065 (0.018) | 0.272 | 3.55 | 0.016 |
| MTG pole ↔︎ Cu | 0.073 | 0.083 (0.023) | 0.271 | 3.53 | 0.017 |
| MTG ↔︎ pMTG | 0.071 | 0.751 (0.212) | 0.267 | 3.48 | 0.020 |
| MTG pole ↔︎ IOG | 0.069 | 0.240 (0.069) | 0.262 | 3.41 | 0.025 |
| ITG ↔︎ Cu | 0.066 | 0.169 (0.050) | 0.257 | 3.34 | 0.032 |
| MTG ↔︎ SOG | 0.063 | 0.211 (0.064) | 0.251 | 3.27 | 0.040 |
| **Noncanonical Sentence Comprehension with Repetition covariate** | | | | | |
| ITG ↔︎ Cu | 0.116 | 0.604 (0.150) | 0.341 | 3.91 | 0.004 |
| MTG pole ↔︎ SOG | 0.106 | 0.217 (0.056) | 0.326 | 3.72 | 0.008 |
| ITG ↔︎ SOG | 0.102 | 0.739 (0.197) | 0.320 | 3.65 | 0.011 |
| ITG ↔︎ pMTG | 0.096 | 2.373 (0.653) | 0.310 | 3.54 | 0.016 |
| MTG pole ↔︎ Cu | 0.092 | 0.228 (0.065) | 0.303 | 3.45 | 0.021 |
| ITG ↔︎ pITG | 0.083 | 1.829 (0.547) | 0.287 | 3.26 | 0.038 |
| STG ↔︎ pSTG | 0.081 | 1.462 (0.443) | 0.284 | 3.22 | 0.043 |

See Figure 3 for region abbreviation definitions. All regions are left hemisphere, unless indicated as right hemisphere with (RH). β_u_ = unstandardized estimated beta coefficient, SE = standard error, β_s_ = standardized estimated beta coefficient, Z = Z-score, BF_m_ (null) = Bayes Factor index indicating support for the null hypothesis.
